# Inhibition of DNMT1 methyltransferase activity via glucose-regulated *O*-GlcNAcylation alters the epigenome

**DOI:** 10.1101/2022.05.11.491514

**Authors:** Heon Shin, Amy Leung, Kevin R. Costello, Parijat Senapati, Hiroyuki Kato, Michael Lee, Dimitri Lin, Xiaofang Tang, Zhen Bouman Chen, Dustin E. Schones

## Abstract

The DNA methyltransferase activity of DNMT1 is vital for genomic maintenance of DNA methylation. We report here that DNMT1 function is regulated by *O*-GlcNAcylation, a protein modification that is sensitive to glucose levels, and that elevated *O*-GlcNAcylation of DNMT1 from high glucose environment leads to alterations to the epigenome. Using mass spectrometry and complementary alanine mutation experiments, we identified S878 as the major residue that is *O*-GlcNAcylated on DNMT1. Functional studies further revealed that *O*-GlcNAcylation of DNMT1-S878 results in an inhibition of methyltransferase activity, resulting in a general loss of DNA methylation that is preferentially at partially methylated domains (PMDs). This loss of methylation corresponds with an increase in DNA damage and apoptosis. These results establish *O*-GlcNAcylation of DNMT1 as a mechanism through which the epigenome is regulated by glucose metabolism and implicates a role for glycosylation of DNMT1 in metabolic diseases characterized by hyperglycemia.

## Introduction

Protein *O*-GlcNAcylation is a dynamic and reversible post-translational modification that attaches a single *O*-linked β-*N*-acetylglucosamine to serine or threonine residues (Hart et al., 1996). It is modulated by two *O*-GlcNAc cycling enzymes, *O*-GlcNAc transferase (OGT) and *O*-GlcNAcase (OGA) that respond to metabolic signals (Hart et al., 2007; Slawson et al., 2010). Increased concentrations of UDP-GlcNAc that are observed in conditions of excess glucose lead to a general increase in protein *O*-GlcNAcylation (Walgren et al., 2003). Obesogenic diets, furthermore, have elevated protein *O*-GlcNAcylation in various human cell types, including liver cells (Guinez et al., 2011), lymphocytes (Torres and Hart, 1984), and immune cells (de Jesus et al., 2018).

As with other post-translational modifications, *O*-GlcNAcylation of proteins can influence the function and/or stability of the targeted proteins (Shin et al., 2018; Yang and Qian, 2017). Thousands of proteins are targets for *O*-GlcNAcylation, including many epigenetic regulatory proteins. For example, the *O*-GlcNAcylation of TET family proteins alter their activity, localization and targeting (Chen et al., 2013; Ito et al., 2014; Shi et al., 2013; Zhang et al., 2014). While all DNA methyltransferases have been shown to be *O*-GlcNAcylated (Boulard et al., 2020), the functional consequences of this have not been previously investigated.

Among the DNA methyltransferase (DNMT) family of proteins, DNMT1 is imperative for maintaining DNA methylation patterns during replication (Bestor and Ingram, 1983). DNMT1 is a modular protein with several domains necessary for interacting with cofactors, including the BAH1 and BAH2 domains (Maresca et al., 2015; Ren et al., 2018). The stability and function of DNMT1 has been shown to be regulated through post-translational modifications, including acetylation, phosphorylation, and methylation (Scott et al., 2014).

Partially methylated domains, large domains with a loss of DNA methylation, were originally identified in cultured cell lines (Lister et al., 2009) and subsequently found to be a characteristic of cancer cells (Berman et al., 2011; Brinkman et al., 2019). PMDs have also been detected in non-cancerous healthy tissues, where they are associated with late replication loci (Hansen et al., 2010; Zhou et al., 2018). While PMDs are generally thought to arise from a lack of fidelity in maintenance methylation (Decato et al., 2020), the mechanisms responsible for the establishment of PMDs have remained unclear. Here, we report that the activity of DNMT1 is regulated by extracellular levels of glucose through *O*-GlcNAcylation, resulting in loss of methylation within PMDs.

## Results

### High glucose conditions increase *O*-GlcNAcylation of DNMT1

To validate that DNMT1 can be *O*-GlcNAcylated, we treated Hep3B cells with OSMI-4 (OSMI), an OGT inhibitor (Martin et al., 2018), as well as with Thiamet-G (TMG), an OGA inhibitor (Elbatrawy et al., 2020). As expected, immunoblots of cellular lysate with an antibody recognizing pan-*O*-GlcNAc (RL2) reveal that inhibition of OGA increased global levels of *O*-GlcNAc while inhibition of OGT decreased global levels of *O*-GlcNAc (Figure 1—figure supplement 1). To distinguish whether DNMT1 is *O*-GlcNAcylated, DNMT1 immunoprecipitation was performed with cellular lysates treated with OSMI or TMG. Immunoblots with *O*-GlcNAc antibodies revealed that TMG treatment increases *O*-GlcNAc of DNMT1 while OSMI treatment decreases *O*-GlcNAc (Figure 1—figure supplement 1). In addition to Hep3B cells, we found that DNMT1 is *O*-GlcNAcylated in HepG2 cells (Figure 1—figure supplement 2) and B cell derived lymphocytes, indicating DNMT1 is *O*-GlcNAcylated across various cell types (Figure 1—figure supplement 2).

**Figure 1.**
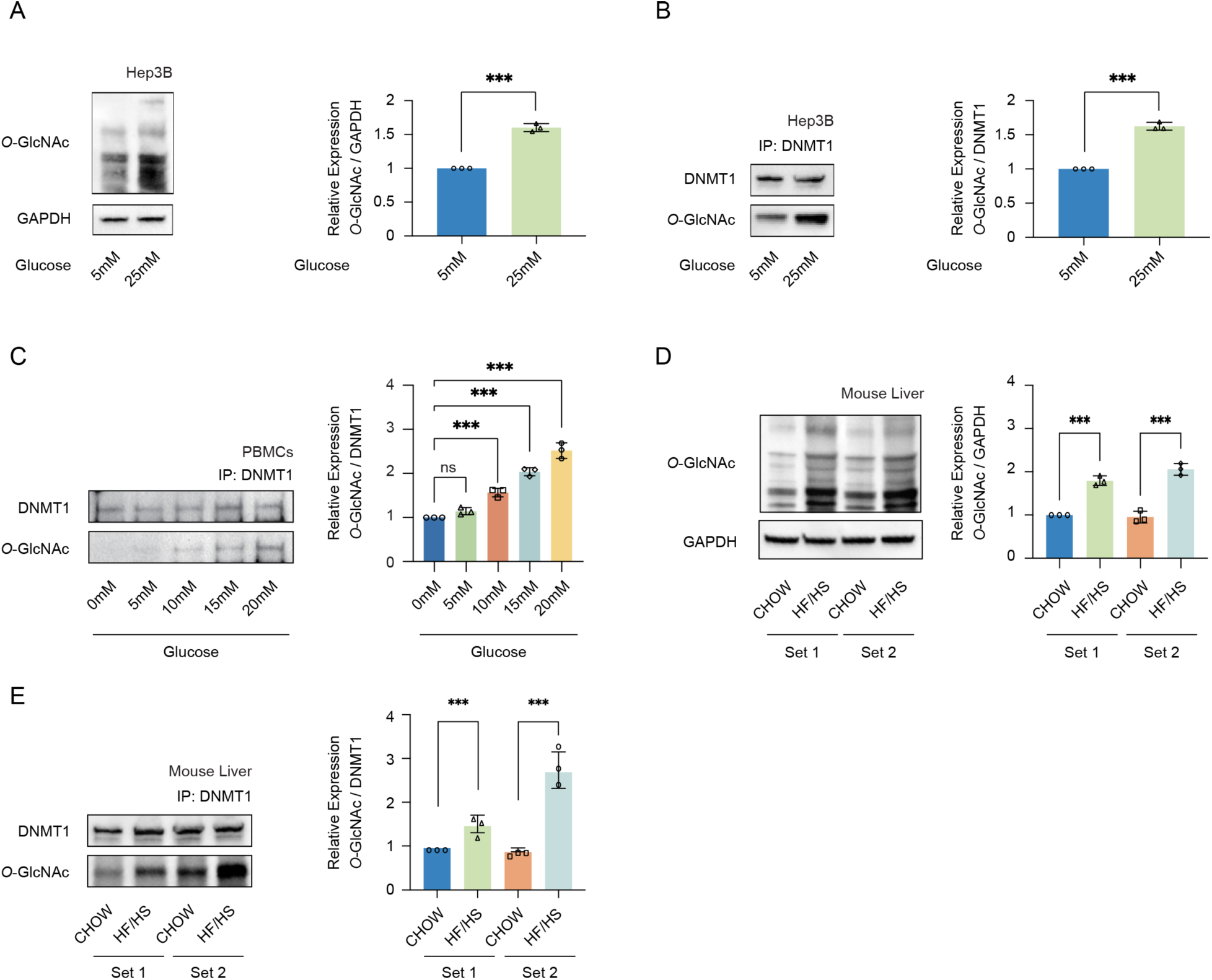
High glucose increases *O*-GlcNAcylation of DNMT1 in cell lines and primary cells. (**A**) Hep3B cells were treated glucose (5mM or 25mM). Shown are immunoblots of collected lysates using antibody targeting *O*-GlcNAc, and GAPDH (*n* = 3). (**B**) Lysates of Hep3B treated with glucose were immunoprecipitated with DNMT1 and immunoprecipitates were immunoblotted with antibody targeting *O*-GlcNAc (*n* = 3). (**C**) PBMCs were isolated from three individual donor blood samples and treated with increasing concentration of glucose for 24 hours. Collected cell lysates from PBMCs were immunoprecipitated with antibody targeting DNMT1 and immunoblotted for *O*-GlcNAc. Representative blot from one donor. (*n* = 3). (**D**) Immunoblots for *O*-GlcNAc, and GAPDH from liver samples of C57BL/6J mice given a high-fat / high-sucrose diet (HF/HS) or normal diet (CHOW) for 4 months. (**E**) Lysates of mouse liver were immunoprecipitated with DNMT1 and immunoprecipitates were immunoblotted with antibody targeting *O*-GlcNAc. \*\*\**p* < 0.0001 by Student’s *t*-test (**A-E**); ns, not significant; Data are represented as mean ± SD from three replicates of each sample. **Figure supplement 1.** DNMT1 can be *O*-GlcNAcylated in Hep3B cells. **Figure supplement 2.** DNMT1 can be *O*-GlcNAcylated in HepG2 cells and B cells derived lymphocytes. **Figure supplement 3.** Global protein *O*-GlcNAcylation was induced with high concentrations of sucrose. **Figure supplement 4.** The enzymatic activity of OGT or OGA was not significantly changed by glucose treatment. **Figure supplement 5.** DNMT1 can be *O*-GlcNAcylated in primary cells (peripheral blood mononuclear cells, PBMCs).

To assess the effect of increased glucose metabolism on *O*-GlcNAcylation of DNMT1, we treated Hep3B cells with normal, or low concentrations of glucose (5 mM) or high glucose (25 mM) and examined global protein *O*-GlcNAcylation as well as the *O*-GlcNAcylation of DNMT1 specifically (Hardiville et al., 2020). Consistent with previous reports (Andrews et al., 2000), the total amount of protein *O*-GlcNAcylation was increased with high glucose treatment (Figure 1A). Global protein *O*-GlcNAcylation was also induced with high concentrations of sucrose, albeit to a lower extent than with glucose (Figure 1—figure supplement 3). To specifically assess the level of *O*-GlcNAcylated DNMT1, we performed immunoprecipitation of DNMT1 from lysates of glucose treated Hep3B cells and immunoblotted for *O*-GlcNAc. As with the analysis of total protein, high glucose treatment increased the *O*-GlcNAcylation of DNMT1 (Figure 1B). High sucrose treatment increased the *O*-GlcNAcylation of DNMT1 as well (Figure 1—figure supplement 3). High glucose and sucrose treatment increased the *O*-GlcNAcylation of DNMT1 in HepG2 cells as well (Figure 1—figure supplement 3). The enzymatic activity of OGT or OGA was not significantly changed by glucose treatment (Figure 1—figure supplement 4), consistent with previous results (Seo et al., 2016).

To examine whether an increase of *O*-GlcNAcylation of DNMT1 also occurs in primary cells, we collected peripheral blood mononuclear cells (PBMCs) from three separate patient donors and treated the PBMCs with increasing glucose levels (0mM, 5mM, 10mM, 15mM, and 20mM). Consistent with our observations in Hep3B cells, we observed an increase in *O*-GlcNAcylation of DNMT1 with increased glucose (Figure 1C). Combining the high glucose condition with OGA inhibition by Thiamet-G (TMG) resulted in a further increase in *O*-GlcNAcylation of DNMT1 (Figure 1—figure supplement 5). To examine the relationship between glucose levels and the *O*-GlcNAcylation of DNMT1 in an *in vivo* context, we examined liver samples from C57BL/6J mice fed an obesogenic high-fat/high-sucrose (HF/HS) diet for 16 weeks (Tang et al., 2020) (details in Methods). These samples displayed an increase in total *O*-GlcNAcylation in liver samples from HF/HS fed mice (Figure 1D) as well as increased *O*-GlcNAcylation of DNMT1 (Figure 1E). These data validate the *O*-GlcNAcylation of DNMT1 and that the degree of *O*-GlcNAcylation of DNMT1 increases with glucose concentrations.

### Identification of the major *O*-GlcNAcylation sites of DNMT1

To begin to identify the major residues *O*-GlcNAcylated on DNMT1, we utilized OGTSite (Kao et al., 2015) to predict potential sites of *O*-GlcNAcylation. OGTSite, which uses experimentally verified *O*-GlcNAcylation sites to build models of substrate motifs, identified 16 candidate *O*-GlcNAc modified sites on human DNMT1 (Table S1). We next employed mass spectrometry analysis to examine the post-translational modifications on DMNT1 in Hep3B cells. We overexpressed DNMT1 using Myc-tagged DNMT1 construct to increase the protein level of DNMT1 in Hep3B cells (Li et al., 2006). Immunoblots with Myc antibody (Yompakdee et al., 1996) revealed a band corresponding to Myc-DNMT1 in transfected, but not mock transfected, cells (Figure 2—figure supplement 1). We further confirmed with immunoprecipitation followed by immunoblot that the overexpressed Myc-DNMT1 can be *O*-GlcNAcylated (Figure 2—figure supplement 1). For mass spectrometry analysis, we treated Myc-DNMT1 expressing cells with 25mM Thiamet-G (TMG) to further increase the *O*-GlcNAcylation of DNMT1. Myc-DNMT1 was enriched from transfected cells by monoclonal Ab-crosslinked immunoprecipitation and subjected to in-solution digestion using three different enzymes (AspN, chymotrypsin, and LysC) and high-resolution LC-MS/MS analysis. Peptide analyses revealed that S878, which is located on the bromo-associated homology (BAH1) domain of DNMT1 is *O*-GlcNAcylated (Figure 2A, B, Figure 2—figure supplement 2, and Table S2). In addition, eight unreported phosphorylated residues were newly detected (T208, S209, S873, S874, S953, S954, S1005, and S1202) (Table S2).

**Figure 2.**
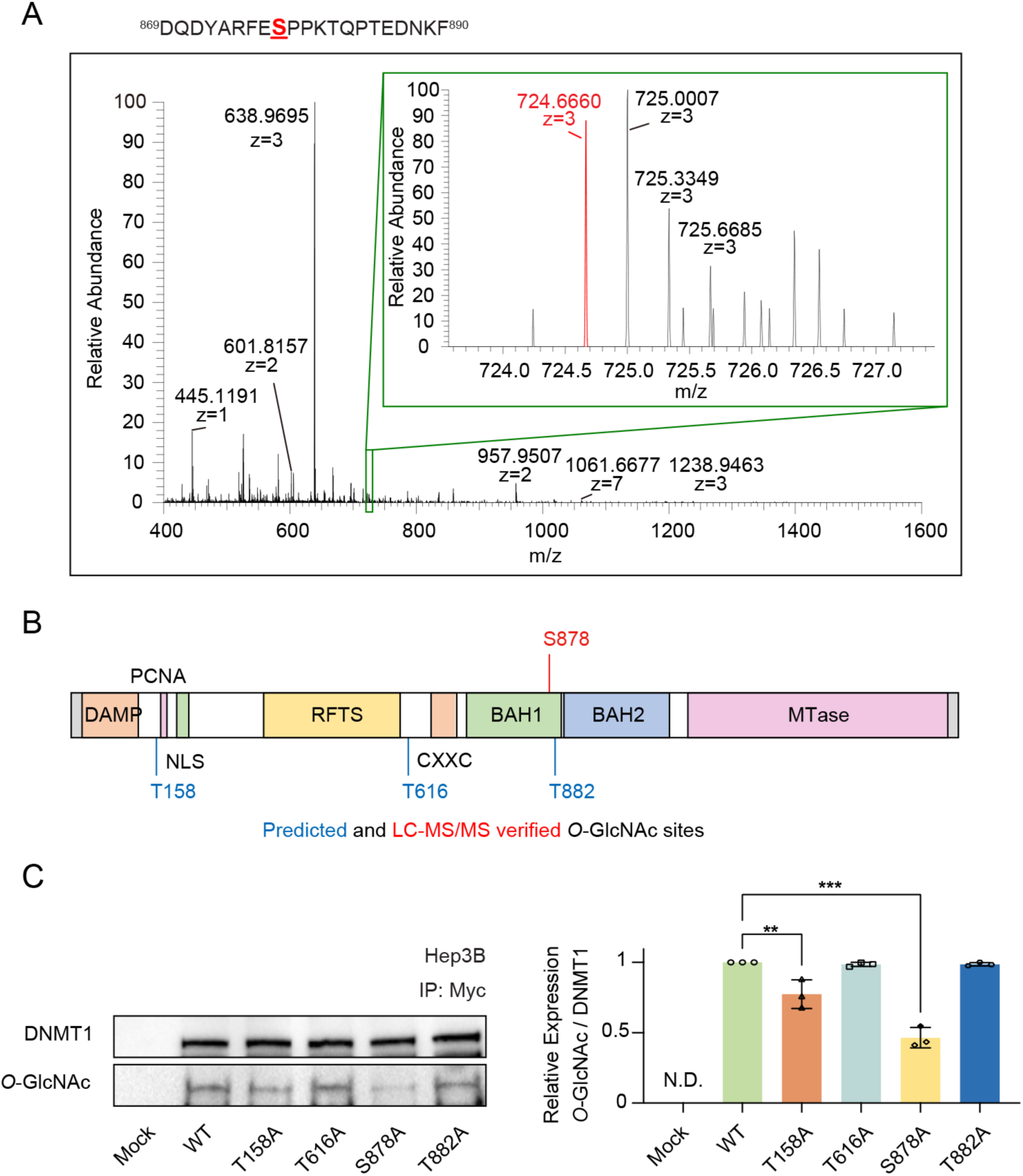
Identification of *O*-GlcNAcylated sites within DNMT1 by LC-MS/MS. (**A**) Schematic drawing of the DNMT1 *O*-GlcNAc modified region enriched from Hep3B cells based on mass spectrometry (MS) data and tandem MS (MS/MS) peaks. FTMS + p NSI Full MS [400.0000-1600.0000]. DQDYARFESPPKTQPTEDNKF (S9 HexNAc) – S878. (**B**) Schematic diagram of identified novel *O*-GlcNAcylated sites within DNMT1 as determined via LC-MS/MS and OGTSite. DMAP, DNA methyltransferase associated protein-binding domain; PCNA, proliferating cell nuclear antigen-binding domain; NLS, nuclear localization sequences; RFTS, replication foci targeting sequence domain; BAH, bromo-adjacent homology domain. (**C**) Each immunoprecipitated Myc-DNMT1 wild type and substituted mutants was immunoblotted with an *O*-GlcNAc antibody (*n* = 3). \*\**p* < 0.0005; \*\*\**p* < 0.0001 by Student’s *t*-test (**C**); N.D., not detected; Data are represented as mean ± SD from three replicates of each sample. **Figure supplement 1.** Myc-DNMT1-WT in Hep3B cells can be *O*-GlcNAcylated. **Figure supplement 2.** Tandem MS/MS peaks of *O*-GlcNAcylated DNMT1 peptides. **Figure supplement 3.** Loss of both threonine and serine in DNMT1 (DNMT1-T158A/S878A) resulted in a loss of *O*-GlcNAcylation.

We chose the three top candidates based on prediction score (T158, T616, and T882) as well as the site identified from mass spectrometry analysis (S878) for further analysis with alanine mutation experiments. The threonine/serine residues were mutated to alanine residues on the Myc-DNMT1 construct and *O*-GlcNAcylation was evaluated with immunoblot following immunoprecipitation. Loss of threonine and serine at positions T158 and S878 respectively resulted in a loss of *O*-GlcNAcylation, indicating that these two residues are required for *O*-GlcNAcylation, with the DNMT1-S878A and DNMT1-T158A/S878A mutant resulting in > 50% reduction of *O*-GlcNAcylation (Figure 2C and Figure 2—figure supplement 3). These results indicate that T158 (near the PCNA binding domain) and S878 (within the BAH1 domain) are the *O*-GlcNAcylated residues of DNMT1.

### *O*-GlcNAcylation of DNMT1 results in loss of DNA methyltransferase activity

The BAH domains of DNMT1 are known to be necessary for DNA methyltransferase activity (Gong et al., 2021; Yarychkivska et al., 2018). Given that S878 is in the BAH1 domain, we reasoned that *O*-GlcNAcylation of this residue could impact the DNA methyltransferase activity of DNMT1. To test this, we treated Hep3B and HepG2 cells with either low (5mM, CTRL) or high glucose combined with TMG (25mM, *O*-GlcNAc) and evaluated the DNA methyltransferase activity of immunoprecipitated DNMT1 with the EpiQuik DNMT Activity/Inhibition ELISA Easy Kit (EpiGentek, details in Methods). Intriguingly, high glucose combined with TMG treatment reduced the activity of DNMT1 (Figure 3A and Figure 3—figure supplement 1).

**Figure 3.**
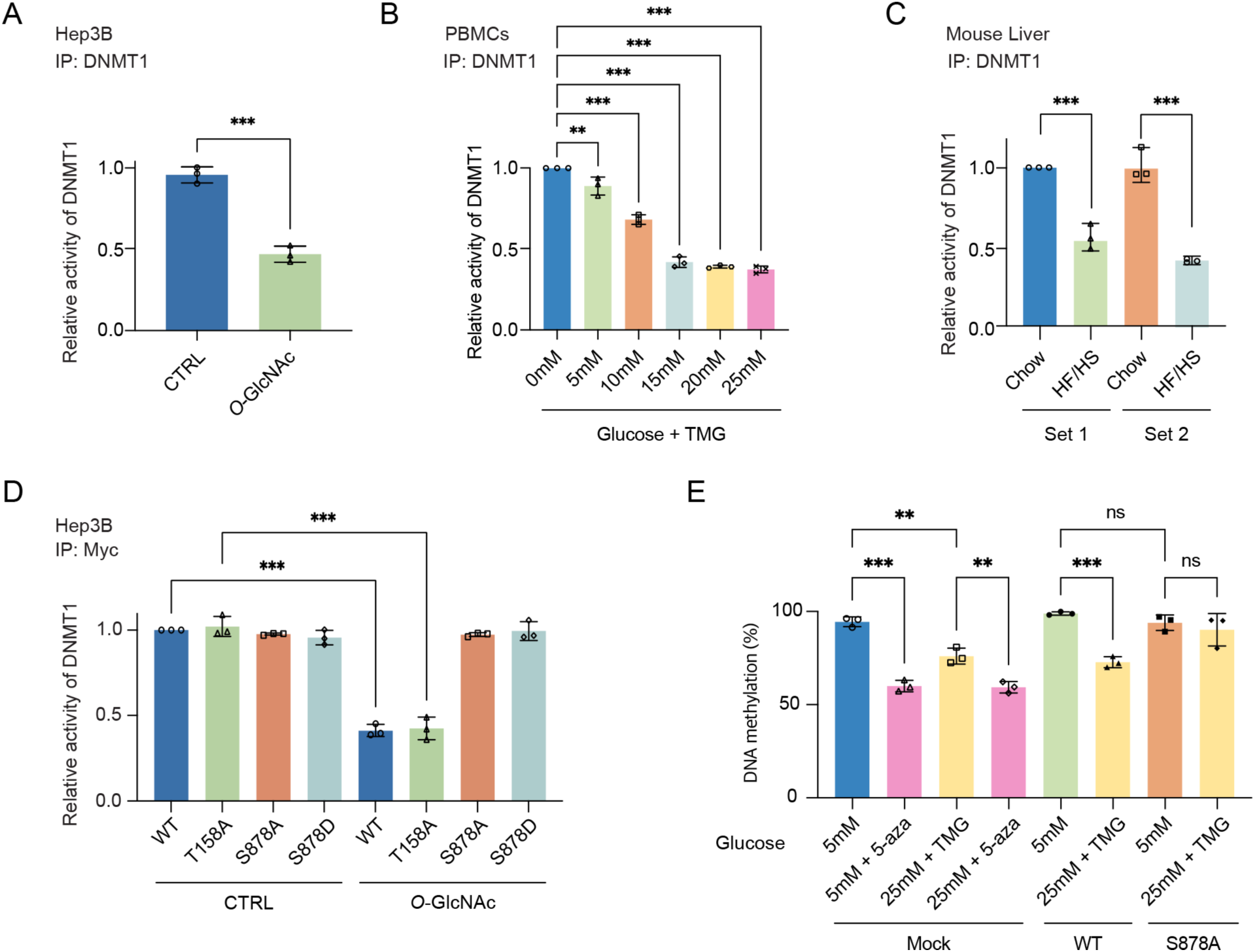
Site specific *O*-GlcNAcylation inhibits DNMT1 methyltransferase function. For (**A-D**), bar graphs are of relative activity of DNA methyltransferase activity measured as absorbance from a DNMT Activity/Inhibition ELISA kit and representative immunoblots of immunoprecipitates performed with antibodies targeting DNMT1. (**A**) Hep3B cells were treated with 5mM (CTRL) or 25mM glucose with TMG (*O*-GlcNAc) (*n* = 3). (**B**) PBMCs from donors were treated with increasing concentrations of glucose (range: 0-25mM with TMG) (*n* = 3). (**C**) Liver samples from C57BL/6J mice given a HF/HS diet or a normal diet (CHOW) for 4 months. (**D**) Immunoprecipitated DNMT1 wild type and substituted mutants were treated with 5mM or 25mM glucose (*n* = 3). (**E**) Each Hep3B and Myc-DNMT1 overexpressed mutant (DNMT1-WT or DNMT1-S878A) was treated with 5mM or 25mM glucose or 5-aza (negative control). Shown are absorbance of global DNA methylation of LINE-1 performed with global DNA methylation LINE-1 kit. (*n* = 3). \*\**p* < 0.005; \*\*\**p* < 0.0001 by Student’s *t*-test (**A-E**); ns, not significant; Data are represented as mean ± SD from three replicates of each sample. **Figure supplement 1.** Site specific *O*-GlcNAcylation at DNMT1 sites abrogate the function of methyltransferase and DNA loss of methylation at CpG island under high glucose/TMG conditions. **Figure supplement 2.** The methylation loss by high glucose/TMG conditions was not apparent in the DNMT1-S878A mutant.

We next examined the impact of glucose levels on the function of DNMT1 in primary cells by treating PBMCs with increasing concentrations of glucose (0mM, 5mM, 10mM, 15mM, 20mM and 25mM with Thiamet-G) for 96 hours and measuring the DNA methyltransferase activity of DNMT1. We observed a striking dose-dependent inhibition of the DNA methyltransferase activity of DNMT1 (Figure 3B). Lastly, we examined the activity of DNMT1 in the liver samples of mice fed a HF/HS diet, which showed a decreased activity of DNMT1 (Figure 3C) compared to chow fed mice. Together, these data indicate that elevated levels of extracellular glucose can inhibit the methyltransferase function of DNMT1.

We next examined the ability of the DNMT1 alanine mutants (DNMT1-T158A and DNMT1-S878A), which cannot be *O*-GlcNAcylated, to attenuate the impact of high glucose- and TMG-induced loss of DNA methyltransferase activity. Compared to the DNA methyltransferase activity of DNMT1-WT (Myc-DNMT1-WT), the DNA methyltransferase activity of DNMT1-S878A (Myc-DNMT1-S878A) is not inhibited by high glucose treatment (Figure 3D), indicating *O*-GlcNAcylation of DNMT1-S878 is directly involved in the inhibition of methyltransferase activity. In contrast, the DNA methyltransferase activity of DNMT1-T158A (Myc-DNMT1-T158A) is inhibited by high glucose combined with TMG treatment in a manner similar to DNMT1-WT (Myc-DNMT1-WT), indicating that *O*-GlcNAcylation of DNMT1-T158 does not affect its DNA methyltransferase activity (Figure 3D).

A previous phospho-proteomic analysis revealed that DNMT1-S878 can be phosphorylated (Zhou et al., 2013), but the functional consequences of this have not been investigated. To evaluate the potential that phosphorylation, rather than *O*-GlcNAcylation, of S878 is leading to the loss of DNA methyltransferase activity, we generated DNMT1-S878D mutant, a phosphomimetic mutant that cannot be *O*-GlcNAcylated and examined DNA methyltransferase activity in normal and high glucose conditions. This phospho-mimetic mutant did not have loss of DNA methyltransferase activity under high glucose conditions, indicating that *O*-GlcNAcylation of S878 but not phosphorylation of S878 is leading to loss of methyltransferase activity of DNMT1 (Figure 3D).

### *O*-GlcNAcylation of DNMT1 results in subsequent loss of DNA methylation

Given our observations that *O*-GlcNAcylation of DNMT1 inhibits its DNA methyltransferase activity, we reasoned that this would further result in a general loss of DNA methylation. To begin to assess this, DNA methylation was assayed using the global DNA methylation LINE-1 kit (Active Motif, details in Methods) as a proxy for global methylation. Comparison of DNA methylation levels under high glucose and TMG with a DNA methylation inhibitor (5-aza; details in Methods) revealed that high glucose leads to a loss of DNA methylation in a manner comparable with the DNA methylation inhibitor (Figure 3E). This methylation loss was not apparent in the DNMT1-S878A mutant, further demonstrating that *O*-GlcNAcylation of S878 within DNMT1 directly affects DNA methylation under high glucose conditions (Figure 3E and Figure 3—figure supplement 2). A complementary assessment of DNA methylation using methylation sensitive restriction enzymes and gel electrophoresis (details in Methods) revealed similar trends (Figure 3—figure supplement 3).

### *O*-GlcNAcylation of DNMT1 results in loss of DNA methylation at partially methylated domains (PMDs)

To more thoroughly examine the impact of high glucose induced *O*-GlcNAcylation of DNMT1 on the epigenome, Myc-DNMT1 overexpressed cell lines (DNMT1-WT and DNMT1-S878A) were treated with either low (5mM, CTRL) or high glucose combined with TMG (25mM, *O*-GlcNAc) and DNA methylation was profiled with nanopore sequencing (ONT PromethION; details in Methods). Comparison of the methylation profiles in these cells revealed a global loss of methylation in high glucose compared to control (Figure 4A, B, and Figure 4—figure supplement 1). Conversely, for the DNMT1-S878A mutant, there was no appreciable decrease in DNA methylation by high glucose (Figure 4A). These results collectively indicate that *O*-GlcNAcylation of S878 of DNMT1 leads to a global loss of DNA methylation.

**Figure 4.**
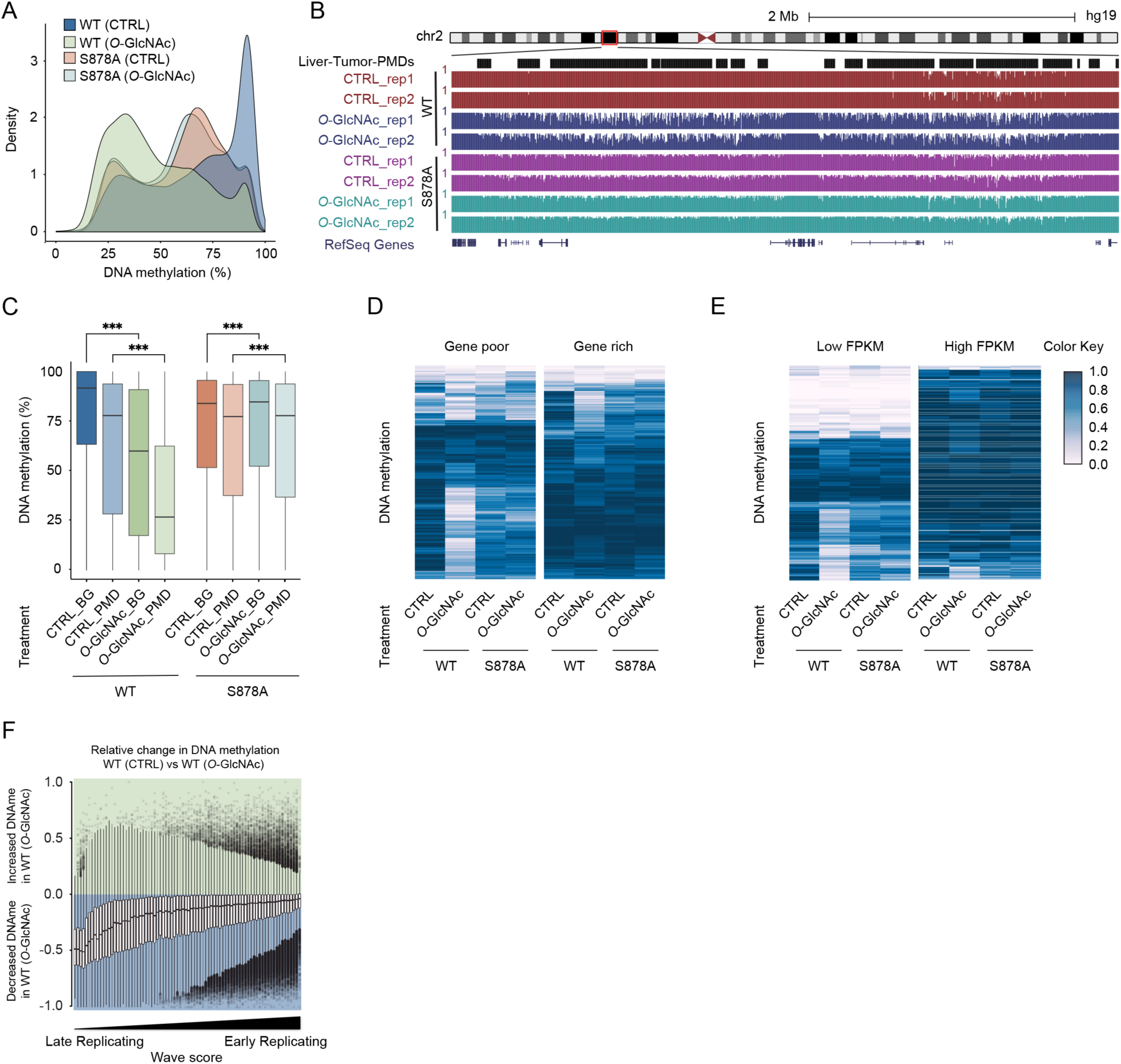
High glucose leads to loss of DNA methylation at cancer specific partially methylated domains (PMDs). (**A**) Density plot of DNA methylation for DNMT1-WT and DNMT1-S878A cells and low (5mM, CTRL) or high glucose with TMG (25mM, *O*-GlcNAc). (**B**) Genome browser screenshot of DNA methylation for DNMT1-WT and DNMT 1-S878A cells and low or high glucose along with liver tumor PMDs from (Li et al., 2016). (**C**) Boxplots of DNA methylation at PMDs or general genomic background (BG) for each DNMT1-WT and DNMT1-S878A treated with low (5mM, CTRL) or high glucose with TMG (25mM, *O*-GlcNAc). (**D**) Heatmap representation of global DNA methylation for DNMT1-WT and DNMT1-S878A cells under low (5mM, CTRL) or high glucose with TMG (25mM, *O*-GlcNAc) at gene poor and gene rich regions. (**E**) Heatmap represent global DNA methylation of wild type and DNMT1 mutants between low FPKM regions and high FPKM regions (DNMT1-WT or DNMT1-S878A) which treated low (5mM, CTRL) or high glucose with TMG (25mM, *O*-GlcNAc) were determined by Nanopolish call methylation. These are defined ‘low FPKM’ as containing less than 25% of RPKM regions per Mb window, and ‘high FPKM’ as containing more than 75% of RPKM regions per Mb window. (**F**) Methylation changes from *O*-GlcNAcylation of DNMT1 by wave score for replication timing (Hansen et al., 2010; Thurman et al., 2007). \*\*\**p* < 0.0001 by Wilcoxon signed-rank test (**C**). **Figure supplement 1.** DNA loss of methylation by increased global *O*-GlcNAcylation decreases. Density plot of DNA methylation for DNMT1-WT and DNMT1-S878A cells and low or high glucose. **Figure supplement 2.** Methylation changes from *O*-GlcNAcylation of DNMT1 in DNMT1-S878A mutant. The loss of methylation was not observed in DNMT1-S878A mutant cells. **Figure supplement 3.** DNA loss of methylation by increased global *O*-GlcNAcylation decreases around the transposable elements (TEs) regions. **Figure supplement 4.** Evolutionarily recent TEs are more likely to lose methylation than older elements in a variety of systems. **Figure supplement 5.** ZFP57 and ZNF605 demonstrate binding to a significant number of LTR12C elements present in liver cancer PMDs. **Figure supplement 6.** Evolutionarily recent elements are less likely to lose methylation induced by *O*-GlcNAcylation of DNMT1. **Figure supplement 7.** Only loci with greater than 5x coverage were retained for analysis, comprising 90% of CpGs in the genome.

Examination of DNA methylation changes induced by *O*-GlcNAcylation of DNMT1 revealed a preferential loss of DNA methylation at liver cancer PMDs (Li et al., 2016) that was not observed in S878A mutant cells (Figure 4B and C). Partially methylated domains (PMDs) have several defining features, including being relatively gene poor and harboring mostly lowly transcribed genes (Decato et al., 2020). We stratified the genome in terms of gene density and transcription rate (see Methods for details) and found that regions that lose methylation in high glucose conditions are largely gene poor (Figure 4D) and contain lowly transcribed genes (Figure 4E) (Chang et al., 2014). PMDs have furthermore been linked to regions of late replication associated with the nuclear lamina (Brinkman et al., 2019). We therefore examined the correlation between loss of methylation caused by high glucose and replication timing (Thurman et al., 2007). In DNMT1-WT cells, late replication domains preferentially lose DNA methylation in high glucose/TMG conditions compared to early replication domains (Figure 4F). This loss of methylation was not observed in DNMT1-S878A mutant cells (Figure 4—figure supplement 2).

### Evolutionarily young transposable elements (TEs) are protected from loss of methylation in high glucose conditions

One of the major functions of DNA methylation in mammalian genomes is the repression of repetitive elements (Edwards et al., 2017). Furthermore, it has been shown that many chromatin proteins involved in the repression of transposable elements (TEs) are capable of being *O*-GlcNAcylated (Boulard et al., 2020). We therefore examined the potential of *O*-GlcNAcylation of DNMT1 to lead to loss of suppression of TEs. We found that high glucose conditions resulted in methylation loss at TEs in a manner similar to the non-repetitive fraction of the genome (Figure 4—figure supplement 3), with a more dramatic loss of methylation at LINEs and LTRs as compared to SINE elements (Figure 4—figure supplement 3). Given that evolutionarily recent TEs are more likely to lose methylation than older elements in a variety of systems (Almeida et al., 2022; Zhou et al., 2020), we examined the methylation status of two younger subfamilies, LTR12C (*Hominoidea*) and HERVH-int (*Catarrhini*) elements (Figure 4—figure supplement 4). While HERVH-int elements show a loss of methylation similar to the rest of the genome, LTR12C elements do not lose methylation in the same manner (Figure 4—figure supplement 4) suggesting they are protected from the loss of methylation. To identify the possible regulatory mechanisms behind the continued maintenance of methylation of LTR12C, an analysis was performed on all LTR12C’s present within hepatic cancer PMDs relative to ChIP-exo data of KRAB-associated zinc-finger proteins, a family of proteins associated with the regulation of transposons. ZFP57 and ZNF605 (on top of the previously defined ZNF676) demonstrate binding to a significant number of LTR12C elements present in liver cancer PMDs (Figure 4—figure supplement 5). Stratifying all TEs by evolutionary age and examining the methylation changes induced by *O*-GlcNAcylation of DNMT1 for each clade revealed that evolutionarily recent elements are less likely to lose methylation (Figure 4—figure supplement 6).

### Methylation changes at promoter regions of apoptosis and oxidative stress response genes upon inhibition of DNMT1

To further examine the impact of the altered epigenome in high glucose conditions, we examined the methylation levels of promoter regions (defined as the 2kb window upstream and downstream of the transcription start site (TSS)) (Figure 4—figure supplement 1 and Figure 5—figure supplement 1). Hypermethylated (gain of methylation) and hypomethylated (loss of methylation) genes were classified as genes with differentially methylated regions overlapping the promoters (Figure 5—figure supplement 1). Pathway analysis (see Methods for details) revealed that genes with hypomethylated promoters are involved in apoptosis and oxidative stress response pathways (Figure 5—figure supplement 2). Examination of apoptosis replated proteins using an apoptosis proteome array revealed that apoptosis agonist proteins (cleaved-caspase3 and phopho-p53 (S15)) are increased and antagonistic proteins (pro-caspase3, surviving, and claspin) are decreased by high glucose treatment (Figure 5—figure supplement 3). Intriguingly, only cIAP-1 protein is increased in the high glucose treated DNMT1-S878A mutant cells (Figure 5—figure supplement 3).

**Figure 5.**
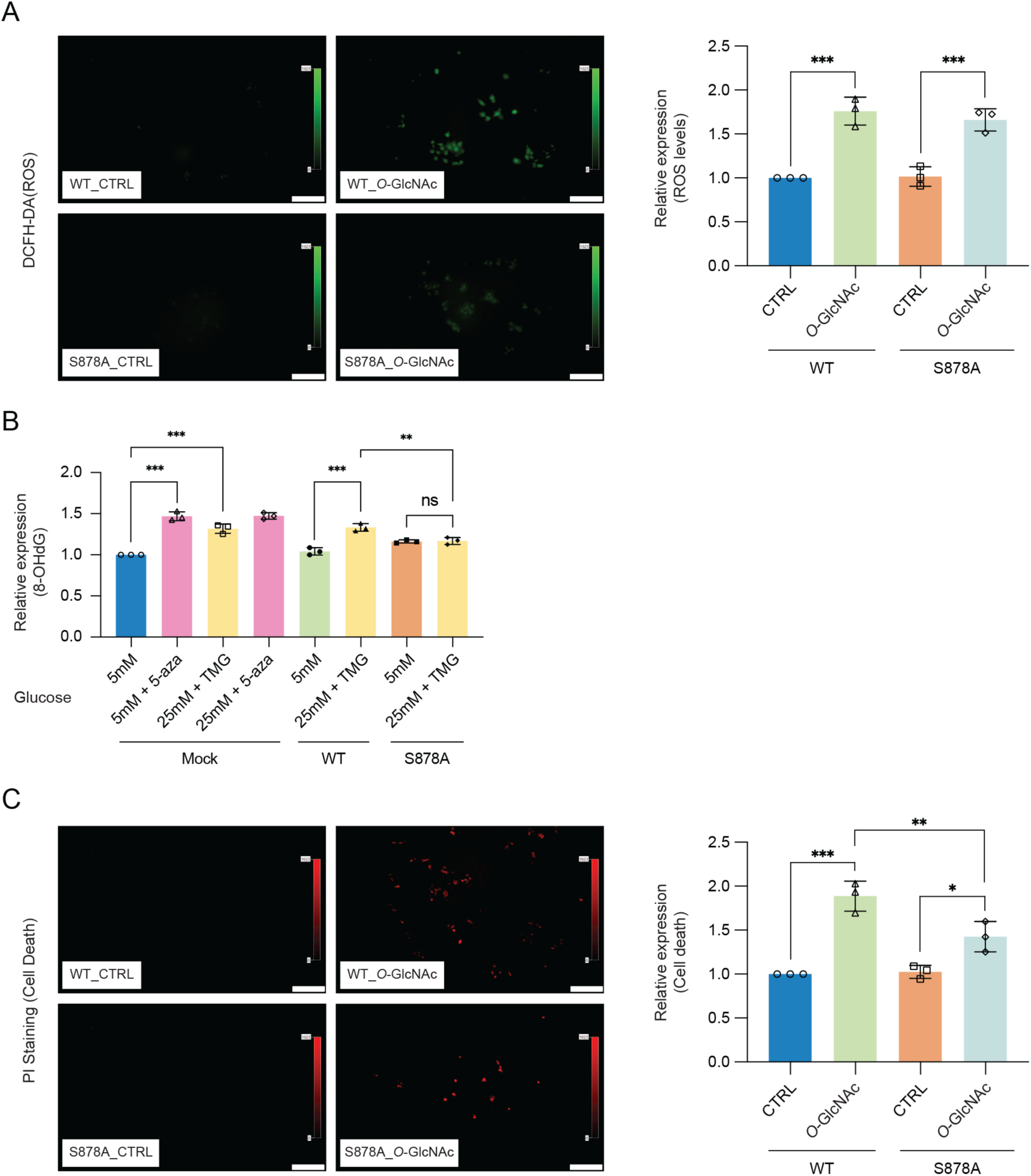
High glucose induced reactive oxygen species (ROS) and DNA damage cause apoptotic cell death in DNMT1-WT cells. (**A**) Quantitative fluorescence image of reactive oxygen species (ROS) in DNMT1-WT and DNMT1-S878A cells and low (5mM, CTRL) or high glucose with TMG (25mM, *O*-GlcNAc). (**B**) Each Hep3B and Myc-DNMT1 overexpressed mutant (DNMT1-WT or DNMT1-S878A) was treated with 5mM or 25mM glucose or 5-aza (negative control). Shown are absorbance of 8-OHdG performed with DNA damage quantification kit. (*n* = 3). (**C**) Quantitative fluorescence image of cell death in propidium iodide staining of DNMT1-WT and DNMT1-S878A cells and low (5mM, CTRL) or high glucose with TMG (25mM, *O*-GlcNAc). \**p* < 0.001; \*\**p* < 0.005; \*\*\**p* < 0.0001 by Student’s *t*-test (**A-C**); ns, not significant; Data are represented as mean ± SD from three replicates of each sample. **Figure supplement 1.** Heatmap representation of promoter DNA methylation for DNMT1-WT and DNMT1-S878A cells under with low (5mM, CTRL) or high glucose with TMG (25mM, *O*-GlcNAc) at gene poor and gene rich regions. **Figure supplement 2.** DNA loss of methylation within promoter region by increased global *O*-GlcNAcylation impact different gene pathways. **Figure supplement 3.** Quantitative analysis of human apoptosis related proteins in DNMT1-WT and DNMT1-S878A by high glucose treatment using Proteome profiler.

### DNA hypomethylation induced DNA damage and triggers apoptosis by high glucose

High glucose induced generation of reactive-oxygen-species (ROS) has been shown to result in increased cell death (Allen et al., 2005). Increased ROS has furthermore been shown to result in upregulation of DNMT1 (He et al., 2012; O’Hagan et al., 2011). To further explore the link between glucose levels, DNMT1 and cell death, we treated DNMT1-WT and DNMT1-S878A cells with either low or high glucose combined with TMG for 96 hours and examined the fluorescence of 2’,7’-dichlorofluorescein diacetate (DCFH-DA) as an indicator for ROS (Figure 5A). The levels of ROS were increased upon treatment with 25 mM glucose with TMG in both of DNMT1-WT and DNMT1-S878A (Figure 5A). Given that high levels of ROS can lead to increased DNA damage and subsequent cell death (Rowe et al., 2008), we analyzed 8-hyrdoxy-2’deoxyguanosine (8-OHdG) as a marker for oxidative DNA damage using EpiQuik 8-OHdG DNA damage quantification kit (EpiGentek, details in Methods). Interestingly, DNA damage was reduced in high glucose treated DNMT1-S878A cells as compared to WT cells (Figure 5B). Furthermore, DNA damage induced by low glucose with 5-aza treatment suggests that DNA hypomethylation is associated with increased DNA damage (Figure 5B) (Palii et al., 2008). Finally, examination of propidium Iodide (PI) levels revealed that cell death was prominently increased in the high glucose treated in DNMT1-WT cells but suppressed in the DNMT1-S878A mutants (Figure 5C). Taken together, these results suggest that ROS-induced DNA damage under hyperglycemic conditions is mitigated by DNA methylation; when DNMT1 function is inhibited via *O*-GlcNAcylation and methylation is lost, ROS-induced DNA damage increases, resulting in apoptosis. These results indicate a that extracellular metabolic stress and cell fate are linked through epigenetic regulation.

## Discussion

Although there is a great deal of evidence regarding the important regulatory role of *O*-GlcNAcylation in gene regulation (Brimble et al., 2010), a direct link with DNA methylation has not previously been established. The maintenance methyltransferase DNMT1 is essential for faithful maintenance of genomic methylation patterns and mutations in DNMT1, particularly in the BAH domains, lead to disruption of DNA methylation (Yarychkivska et al., 2018). While it has been shown that DNTM1 can be *O*-GlcNAcylated (Boulard et al., 2020), the site of *O*-GlcNAc modification on DNMT1 as well as the functional consequences of this modification have not previously been examined.

We reveal that *O*-GlcNAcylation of DNMT1 impacts its DNA methyltransferase activity and affects DNMT1 function leading to loss of DNA methylation at partially methylated domains (PMDs). PMDs are observed in both healthy and cancerous cells and have been suggested to be associated with mitotic dysfunction. However, models for how these domains are established remain incomplete (Decato et al., 2020). The results presented here suggest an additional layer whereby *O*-GlcNAcylation of DNMT1 at S878 due to increased glucose levels can inhibit the function of DNA methyltransferase activity of DNMT1, resulting in loss of methylation and establishment of partially methylated domains.

High glucose conditions have previously been reported to lead to an increase in nuclear 25-Hydroxycholesterol, which induces lipid accumulation and activates DNMT1 (Allen et al., 2005; Wang et al., 2020). Our results are consistent with the activity of DNMT1 gradually increased by glucose concentrations (Figure 3—figure supplement 1). This trend is reversed, however, upon TMG treatment (Figure 3—figure supplement 1), suggesting that the increased activity of DNMT1 associated with glucose treatment is directly inhibited by *O*-GlcNAcylation within DNMT1.

Metabolic diseases such as obesity and diabetes have been linked to epigenetic changes that alter gene regulation (Ling and Ronn, 2019). It has previously been established that there is a general increase in protein *O*-GlcNAcylation in hyperglycemia conditions (Vasconcelos-Dos-Santos et al., 2018) and several epigenetic regulatory factors have been shown to have increased *O*-GlcNAcylation under high glucose conditions (Bauer et al., 2015; Etchegaray and Mostoslavsky, 2016; Yang et al., 2020). Our findings that extracellular glucose promotes *O*-GlcNAcylation of DNMT1 and inhibition of DNMT1’s function in maintenance of genomic methylation provide direct evidence that extracellular levels of glucose is linked with epigenomic regulation.

## Materials and Methods

### Antibodies and Regents

Information on antibodies and reagents used in this study are provided in Table S3.

### Cell culture and plasmid DNA transfection

Human hepatocellular carcinoma cell lines HepG2 (HB-8065), and Hep3B (HB-8064) were purchased from ATCC (Manassas, VA, USA). All cell lines were shown to be negative in mycoplasma test using MycoScope (MY01050, Genlantis, San Diego, CA, USA). The following ATCC-specified cell culture media were used: Dulbecco’s modified Eagle’s medium (DMEM, 11885-084, Grand Island, NY, USA) or high glucose Dulbecco’s modified Eagle’s medium (DMEM, 11995-065, Gibco, Grand Island, NY, USA) with 10% fetal bovine serum (FBS, SH30910.03, HyClone, South Logan, UT, USA) and Opti-MEM (1869048, Gibco, Grand Island, NY, USA). All cells were cultured in a 37°C with a 5% CO_2_ atmosphere incubator. HepG2 and Hep3B cells were transiently transfected (with the pcDNA3 with or without DNMT1 cDNA) using Lipofectamine 3000 (Invitrogen, Carlsbad, CA, USA) and selected with Geneticin (G418, 10131-035, Gibco, Grand Island, NY, USA) according to the manufacturer’s instructions. Human DNMT1 plasmid was purchased from Addgene (#36939, Watertown, MA, USA) (Li et al., 2006).

### Isolation of peripheral blood mononuclear cells (PBMCs)

Blood samples from de-identified healthy donors were obtained following guidelines at the City of Hope as describe (Leung et al., 2018). PBMCs such as lymphocyte, monocyte or a macrophage were isolated directly from human whole blood using Ficoll-Paque (Premium, GE Healthcare, Chicago, IL, USA) density gradient centrifugation. 15ml whole blood was mixed with same volume of phosphate-buffered saline containing 0.1% Fetal bovine serum + 2mM EDTA (PBS solution). Next, the blood mix was placed on top of 15 ml Ficoll and centrifuged at 400g to 200g for 40 min without brake. Next, remove the supernatant and washed three times with PBS solution.

### Isolation of B cells and Epstein-Barr virus (EBV) infection for lymphocyte transformation

CD19^+^ B cells were isolated from PBMCs (peripheral blood mononuclear cells) using Dynabeads CD19^+^ pan B (11143D, Invitrogen, Carlsbad, CA, USA) according to the manufacturer’s instruction. 2.5 × 10^8^ cells of PBMCs were resuspended in 10ml Isolation buffer (PBS, 0.1% BSA, 2mM EDTA). 250 μl of pre-washed beads were added to PBMCs and incubated for 20 min in 4°C with gentle rotation. For positive isolation of CD19^+^ B cells, beads and supernatant were separated using magnet, and supernatant was discarded. Beads were washed three times, and beads bounded with CD19^+^ B cells were resuspended with 2.5 ml of cell culture medium (80% RPMI1640, 20 % heat-inactivated FBS, Glutamine). CD19^+^ B cells were released from Dynabeads using DETACHaBEAD (Invitrogen, ca12506D) according to the manufacturer’s instruction.

B cells were infected with Epstein-Barr virus (EBV) to transform lymphocyte. 10 ml of B cells were transferred into T75 flask. 1.5ml of stock EBV collected from a B95-8 strain-containing marmoset cell lines and 1ml of Phytohemagglutinin P (PHA-P) were added to flask and incubated in a 37°C with a 5% CO_2_ atmosphere incubator. Every 5 to 7 days, 10 ml of cell culture medium was added. Cells were let to grow in CO_2_ atmosphere incubator for 30 days until all B cells were transformed to LCL.

### Mouse liver samples

All animal experiments conducted have been approved by the Institutional Animal Care and Use Committees at City of Hope. All of the animals were handled according to approved institutional animal care and use committee (IACUC) protocols (#17010). C57BL/6J mice were randomized to receive irradiated high-fat / high-sucrose (HF/HS) diet (D12266Bi, Research Diets Inc, 17% kcal protein, 32% kcal fat, 51% kcal carbohydrate) starting at 8 weeks old for 16 weeks. Mice on chow diet (D12489Bi, Research Diets Inc, 16.4% kcal protein, 70.8% kcal carbohydrate, 4.6% kcal fat) were fed for the same duration. To reduce blood contamination, mice were washed 10x with phosphate-buffered saline (PBS) solution. Washed liver tissues (two CHOW and two HF/HS) were cut into several pieces and divided into three groups each (three sets per each condition, total 12 samples). Each group of washed liver tissues were lysed with non-detergent IP buffer in the presence of a protease inhibitor (Cat#8340; Sigma-Aldrich) and a phosphatase inhibitor cocktail (Cat# 5870; Cell Signaling) for the western blotting or immunoprecipitation. An increase in fasting blood glucose levels due to the HF/HS diet has been previously reported (Franson et al., 2021; Tang et al., 2020). At the end points, mice were euthanized with CO2 inhalation.

### Immunoprecipitation and western blot analysis

Cell lysates were incubated with specific antibodies and lysis buffer for 4 hours. Subsequently, 30 μl of washed Dynabeads (14311D, Thermo Fisher, Waltham, MA, USA) were added to each lysate and incubated overnight at 4°C. Next, the beads were washed five times, and the antigens were eluted twice using 8M Urea buffer (8M Urea, 20mM Tris pH 7.5, and 100mM NaCl) and concentrated. The resulting samples were separated by Mini-PROTEAN TGX (4-20%, 4561093, Bio-Rad Laboratories, Hercules, CA, USA) and transferred onto nitrocellulose membranes (Amersham Hybond, 10600021, GE Healthcare, Chicago, IL, USA) using Trans-Blot SD Semi-dry Transfer Cell system (Bio-Rad Laboratories, Hercules, CA, USA). The membranes were then blocked with 5% skim milk in Tris-buffered saline + Tween-20 (TBS-T; 20 mM Tris, 137 mM NaCl, 0.1% Tween-20, pH 7.6), incubated overnight at 4°C with a 1:1000 dilution of each antibodies, and subsequently incubated for 1 h with a 1:5000 dilution of a horseradish peroxidase-conjugated goat anti-mouse secondary antibody (ab6789, Abcam, Cambridge, UK) or goat anti-rabbit secondary antibody (ab6721, Abcam, Cambridge, UK). Immunoreactive proteins were detected using SuperSignal West Dura Extended Duration Substrate (34076, Thermo, Rockford, IL, USA) and detected using a ChemiDoc MP Imaging system (Bio-Rad Laboratories, Hercules, CA, USA). The band intensity was densitometrically evaluated using Image Lab software (Version 5.2, Bio-Rad Laboratories, Hercules, CA, USA).

### Protein identification using the Thermo Fusion Lumos system LC-MS/MS

LC-MS separation was performed on an Thermo Fusion Lumos system (Thermo Scientific, Waltham, MA, USA). For LC separation, 60-minute LC gradient on EasySpray column (particle sizes: 500 mm, 75 μm) was used for peptide separation. Mass spectra for peptide identification or quantification were acquired using an Orbitrap Lumos mass at 120,000 resolutions. Full MS scan ranges were acquired from 156 to 2000 m/z. MS/MS spectra were acquired at a resolution of 30,000 using HCD at 35% collision energy.

Identify post-translational modifications on DNMT1 using complementary in-solution digestion with three enzymes (chymotrypsin, AspN and LysC). Protein was reduced with DTT and alkylated with iodoacetamide. Aliquots of protein were separately digested with three enzymes and peptides were desalted using a C18 tip.

Raw data files were submitted to Byonic (v2.16.11) for target decoy search against the human protein database (uniprot/swissprot, 2020). Peptide-level confidence threshold was set at 99% (FDR <0.01). The sample was bracketed by *E. coli* QC runs, which were then correlated to ensure instrument quality. QC passed threshold (≥ 0.9) with an R^2^ of 0.98 (correlation value, R=0.99)

### Site-directed point mutation

Specific primers for serine (S) and threonine (T) to alanine (A) and aspartic acid (D) mutations of DNMT1 were designed and used to site-directed point mutations in a plasmid vector. A recombinant DNA pcDNA3/Myc-DNMT1 was a gift from Arthur Riggs (Addgene plasmid #36939 Watertown, MA, USA) (Li et al., 2006). A PCR-amplified DNA fragment of pcDNA3-DNMT1 was generated using Q5 Site-Directed Mutagenesis Kit (E0554S, NEB, Ipswich, MA, USA). The primers used in this process are described in Supporting Information. After PCR, the non-mutated sequences were cleaved using Q5 KLD enzyme (New England Biolabs, Ipswich, MA, USA) according to the manufacturer’s instructions. The mutated vectors were transformed into *E. coli* competent cells (NEB 5-alpha, New England Biolabs, Ipswich, MA, USA) that were cultured and prepared using a GenElute HP Plasmid Midi kit (NA0200-1KT, Sigma-Aldrich, St. Louis, MO).

### DNA methyltransferase activity assay

DNA methyltransferase activities of endogenous DNMT1 and recombinant DNMT1 were measured by EpiQuik DNMT Activity/Inhibition ELISA Easy Kit (P-3139-48, EpiGentek, Farmingdale, NY, USA) according to the manufacturer’s instructions. The endogenous DNMT1 (by anti-DNMT1 Ab, 60B1220.1) and recombinant DNMT1 (by anti-Myc, ab18185) were enriched using immunoprecipitation from each cell or tissue lysates. DNMT1s were isolated and normalized by BCA analysis. The activity of 5 ng DNMT1 was analyzed by 450nm ELISA with an optimal wavelength of 655nm.

### Global DNA Methylation LINE-1 assay

Global DNA methylation LINE-1 were measured by Active Motif (55017, Carlsbad, CA, USA) according to the manufacturer’s instructions. Each Hep3B and myc-DNMT1 overexpressed mutants (DNMT1-WT or DNMT1-S878A) were treated 5mM glucose, or 25mM glucose, and 5-aza (negative control). The activity of 100 ng was analyzed by 450nm ELISA with an optimal wavelength of 655nm.

### Agilent 4200 TapeStation

Global DNA methylation were measured by Agilent 4200 TapeStation system (Santa Clara, CA, USA) with the Genomic DNA ScreenTape (5064-5365) and Genomic DNA Reagent (5067-5366) according to the manufacturer’s instructions.

### Nanopore PromethION sequencing

Genomic DNA was isolated from each DNMT1-WT or DNMT1-S878A treated 5mM glucose or 25mM glucose combined with TMG using the QIAGEN DNA Mini Kit (13323, Qiagen) with Genomic-tip (10223, Qiagen) according to the manufacturer’s instructions. Sequencing libraries were prepared using Ligation sequencing kit (SQK-LSK109, Oxford Nanopore Technologies) according to the manufacturer’s instructions. Sequencing was performed on a PromethION (Oxford Nanopore Technologies). Data indexing were performed using nanopolish (RRID:SCR_016157). Reads were aligned using minimap2 (RRID:SCR_018550) with the options -a -x map-ont (Li, 2018). Methylation state of CpGs was called using nanopolish (RRID:SCR_016157) with the options call-methylation -t 8 (Loman et al., 2015). Only loci with greater than 5x coverage were retained for analysis, comprising 90% of CpGs in the genome (Figure 4—figure supplement7). Methylation percentage was averaged across CpG islands.

### Determination of gene-poor or gene-rich regions and FPKM

Hep3B RNA-seq data were obtained from a previous publication (Chang et al., 2014). Fastq files were aligned using HISAT2 version 2.1.0 (RRID:SCR_015530) to the hg19 genome. Duplications are removed using picard version 2.10.1 (RRID:SCR_006525). Aligned reads were sorted using samtools version 1.10 (RRID:SCR_002105). UCSC genome browser tracks were established using bedGraphToBigWig. FPKM was calculated using StringTie version 1.3.4d (RRID:SCR_016323).

For all datasets, Bedgraph files were generated using bedtools version 2.29.0 (RRID:SCR_006646). BigWigs were generated using the UCSCtools bedGraphToBigWig. Heatmap of global DNA methylation for DNMT1-WT and DNMT1-S878A cells under low or high glucose were generated using a custom script to profile the read coverage at each base and were visualized using pheatmap version 1.0.12 (RRID:SCR_016418). All other heatmaps and aggregate plots of loci that extend were generated using deeptools (RRID:SCR_016366).

### Measurement of Reactive Oxygen Species (ROS) and Propidium Iodide (PI) staining

DNMT1-WT and DNMT1-S878A cells were seeded into 12-well plates with 5mM or 25mM glucose and TMG. The medium in each well was replaced with HBSS with 10 μM DCF-DA (2’,7’-dichlorofluorescein diacetate; D6883, Abcam) or propidium iodide staining solution (P4864, Abcam). The fluorescence was filtered with fluorescein isothiocyanate (FITC) for ROS or Texas Red for PI staining (Shin et al., 2020). Averages of fluorescence were analyzed by Olympus Cellsens software.

### DNA damage analysis

DNA damage was analyzed using EpiQuik 8-OHdG DNA Damage Quantification Direct Kit (P-6003, EpiGentek) according to the manufacturer’s protocol. The activity of 200 ng DNA was analyzed by 450nm absorbance microplate reader.

### Apoptosis array analysis

Apoptosis related proteins were analyzed using Proteome profiler human apoptosis array kit (ARY009, R&D Systems) according to the manufacturer’s protocol. The spots were detected using a ChemiDoc MP Imaging system (Bio-Rad Laboratories, Hercules, CA, USA) (Na et al., 2020). The band intensity was densitometrically evaluated using Image Lab software (Version 5.2, Bio-Rad Laboratories, Hercules, CA, USA).

### KZFP binding in PMD-associated LTR12Cs

A list of partially methylated domains (PMD) in Hep3B cells were obtained from a previous publication (Li et al., 2016). A full list of LTR12Cs was generated from filtering hg19-repmask.bed. The PMD-associated LTR12Cs were found using bedtools version 2.29.0 (RRID:SCR_006646). Putative KZFP regulators of LTR12Cs were determined using the consensus sequence of LTR12Cs and ChIP-exo from the Imbeault and Trono studies on the UCSC Repeat Browser (Imbeault et al., 2017). PMD-associated LTR12Cs were then aligned again with peak files containing the ChIP-exo data to acquire a list of PMD-associated LTR12Cs with KZFP binding. Significant binding was defined as > 5 sequences bound, as most LTR12Cs demonstrated very minimal KZFP binding (< 1 sequences bound).

### Quantification and Statistical Analysis

Statistical analyses were performed and graphed using GraphPad Prism 9 (v9.3.1). All statistical tests were performed by three independent experiments assay, and the data are presented as means ± standard deviations. \**p* < 0.001; \*\**p* < 0.0005; \*\*\**p* < 0.0001 by Student’s *t*-test; ns, not significant; Data are represented as mean ± SD.

## Supporting information

Supplemental_informations

## Acknowledgements

This manuscript is dedicated to the memory of Dr. Arthur Riggs. This work was supported by the National Institutes of Health, grants R01DK112041 and R01CA220693 (D.E.S.). Research reported in this publication included work performed in the Pathology and Integrative Genomics Cores of the City of Hope and was supported by the City of Hope CCSG Pilot award from National Cancer Institute of the National Institutes of Health under award number P30CA033572.

## Data availability

PromethION sequencing data have been deposited in the NCBI Gene Expression Omnibus (GEO) and Sequence Read Archive (SRA) under accession no. GSE201470.

## Author Contributions

Heon Shin, Conceptualization, Data curation, Formal analysis, Investigation, Methodology, Validation, Visualization, Writing – original draft, Writing – review and editing; Amy Leung, Conceptualization, Formal analysis, Methodology; Kevin R. Costello, Data curation, Formal analysis; Parijat Senapati, Hiroyuki Kato, Michael Lee, Dimitri Lin, Formal analysis; Xiaofang Tang, Formal analysis, Provide samples; Zhen Bouman Chen, Provide samples; Dustin E. Schones, Conceptualization, Funding acquisition, Investigation, Methodology, Project administration, Supervision, Writing – original draft, Writing – review and editing.

