## Supplemental_informations for "Inhibition of DNMT1 methyltransferase activity via glucose-regulated *O*-GlcNAcylation alters the epigenome"

This PDF file includes:

**Figure 1—figure supplement 1.** DNMT1 can be *O*-GlcNAcylated in Hep3B cells.

**Figure 1—figure supplement 2.** DNMT1 can be *O*-GlcNAcylated in HepG2 cells and B cells derived lymphocytes.

**Figure 1—figure supplement 3.** Global protein *O*-GlcNAcylation was induced with high concentrations of sucrose.

**Figure 1—figure supplement 4.** The enzymatic activity of OGT or OGA was not significantly changed by glucose treatment.

**Figure 1—figure supplement 5.** DNMT1 can be *O*-GlcNAcylated in primary cells (peripheral blood mononuclear cells, PBMCs).

**Figure 2—figure supplement 1.** Myc-DNMT1-WT in Hep3B cells can be *O*-GlcNAcylated.

**Figure 2—figure supplement 2.** Tandem MS/MS peaks of *O*-GlcNAcylated DNMT1 peptides.

**Figure 2—figure supplement 3.** Loss of both threonine and serine in DNMT1 (T158A/S878A) resulted in a loss of *O*-GlcNAcylation.

**Figure 3—figure supplement 1.** Site specific *O*-GlcNAcylation at DNMT1 sites abrogate the function of methyltransferase and DNA loss of methylation at CpG island under high glucose/TMG conditions.

**Figure 3—figure supplement 2.** The methylation loss by high glucose/TMG conditions was not apparent in the DNMT1-S878A mutant.

**Figure 4—figure supplement 1.** DNA loss of methylation by increased global *O*-GlcNAcylation decreases. Density plot of DNA methylation for DNMT1-WT and DNMT1-S878A cells and low or high glucose.

**Figure 4—figure supplement 2.** Methylation changes from *O*-GlcNAcylation of DNMT1 in DNMT1-S878A mutant. The loss of methylation was not observed in DNMT1-S878A mutant cells.

**Figure 4—figure supplement 3.** DNA loss of methylation by increased global *O*-GlcNAcylation decreases around the transposable elements (TEs) regions.

**Figure 4—figure supplement 4.** Evolutionarily recent TEs are more likely to lose methylation than older elements in a variety of systems.

**Figure 4—figure supplement 5.** ZFP57 and ZNF605 demonstrate binding to a significant number of LTR12C elements present in liver cancer PMDs.

**Figure 4—figure supplement 6.** Evolutionarily recent elements are less likely to lose methylation induced by *O*-GlcNAcylation of DNMT1.

**Figure 4—figure supplement 7.** Only loci with greater than 5x coverage were retained for analysis, comprising 90% of CpGs in the genome.

**Figure 5—figure supplement 1.** Heatmap representation of promoter DNA methylation for DNMT1-WT and DNMT1-S878A cells under with low (5mM, CTRL) or high glucose with TMG (25mM, *O*-GlcNAc) at gene poor and gene rich regions.

**Figure 5—figure supplement 2.** DNA loss of methylation within promoter region by increased global *O*-GlcNAcylation impact different gene pathways.

**Figure 5—figure supplement 3.** Quantitative analysis of human apoptosis related proteins in DNMT1-WT and DNMT1-S878A by high glucose treatment using Proteome profiler.

**Table S1.** Prediction of *O*-GlcNAcylated sites within DNMT1 using OGTSite.

**Table S2.** List of identified proteins.

**Table S3.** List of antibodies and reagent used in this study.

**Legends for Figure 1—source data 1.**

**Legends for Figure 1—figure supplement 1-source data 1.**

**Legends for Figure 1—figure supplement 2-source data 1.**

**Legends for Figure 1—figure supplement 3-source data 1.**

**Legends for Figure 1—figure supplement 5-source data 1.**

**Legends for Figure 2—source data 1.**

**Legends for Figure 2—figure supplement 1-source data 1.**

**Legends for Figure 2—figure supplement 3-source data 1.**

**Legends for Figure 5—source data 1.**

**Legends for Figure 5—figure supplement 3-source data 1**

Other supplementary materials for this manuscript include the following:

**Figure 1—source data 1.** Uncropped blot files of Figure 1A-E.

**Figure 1—figure supplement 1-source data 1.** Uncropped blot files of Figure 1—figure supplement 1A and B.

**Figure 1—figure supplement 2-source data 1.** Uncropped blot files of Figure 1—figure supplement 2A and B.

**Figure 1—figure supplement 3-source data 1.** Uncropped blot files of Figure 1—figure supplement 3A-C.

**Figure 1—figure supplement 5-source data 1.** Uncropped blot files of Figure 1—figure supplement 5.

**Figure 2—source data 1.** Uncropped blot files of Figure 2C.

**Figure 2—figure supplement 1-source data 1.** Figure 2—figure supplement 1A and B.

**Figure 2—figure supplement 3-source data 1.** Figure 2—figure supplement 3.

**Figure 5—source data 1.** Raw fluorescence image files of Figure 5A and C.

**Figure 5—figure supplement 3-source data 1.** Uncropped blot files of Figure 5—figure supplement 3.

### Figure supplements

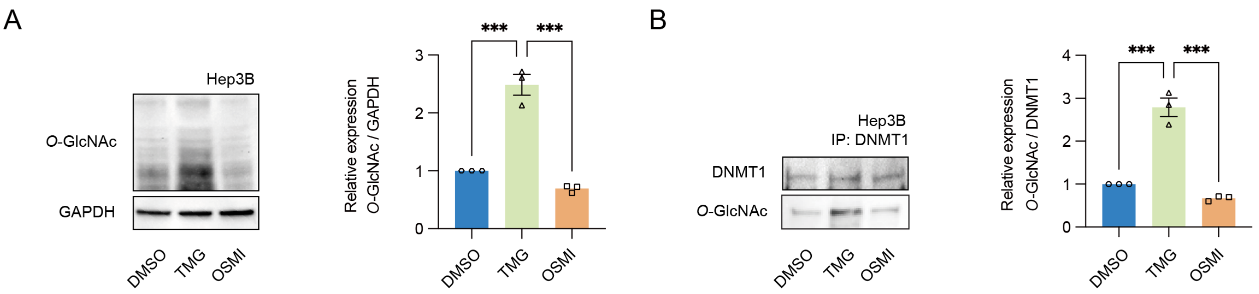

**Figure 1—figure supplement 1.** DNMT1 can be *O*-GlcNAcylated in Hep3B cells. (**A**) Hep3B cells were treated with Thiamet-G (TMG) or OSMI-4 (OSMI). Shown are representative immunoblots of treated Hep3B lysates performed with antibodies targeting *O*-GlcNAc, and GAPDH and bar graphs of relative expression between *O*-GlcNAc compared to control, GAPDH ($n$ = *3*, experimental replicates). (**B**) Lysates from treated Hep3B with glucose were immunoprecipitated with DNMT1 and immunoprecipitates were immunoblotted with antibody targeting *O*-GlcNAc ($n$ = *3*). ***$p < 0.0001$ by Student's $t$-test (**A** and **B**); Data are represented as mean ± SD from three replicates of each sample.

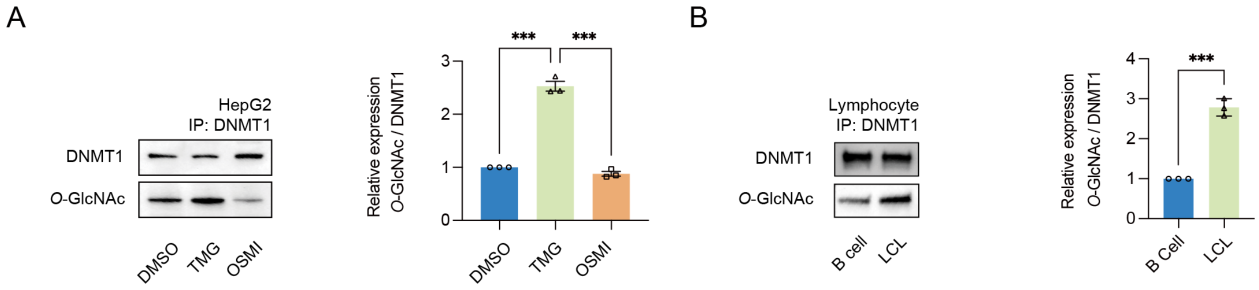

**Figure 1—figure supplement 2.** DNMT1 can be *O*-GlcNAcylated in HepG2 cells and B cells derived lymphocytes. (**A**) HepG2 cells were treated with Thiamet-G or OSMI. Shown are immunoblots of treated HepG2 lysates performed with immunoblots of immunoprecipitates performed with antibodies targeting *O*-GlcNAc ($n = 3$). (**B**) Shown are immunoblots of B cell and lymphocytes (LCL) lysates performed with immunoblots of immunoprecipitates performed with antibodies targeting *O*-GlcNAc ($n = 3$). ***$p < 0.0001$ by Student's *t*-test (**A** and **B**); Data are represented as mean ± SD from three replicates of each sample.

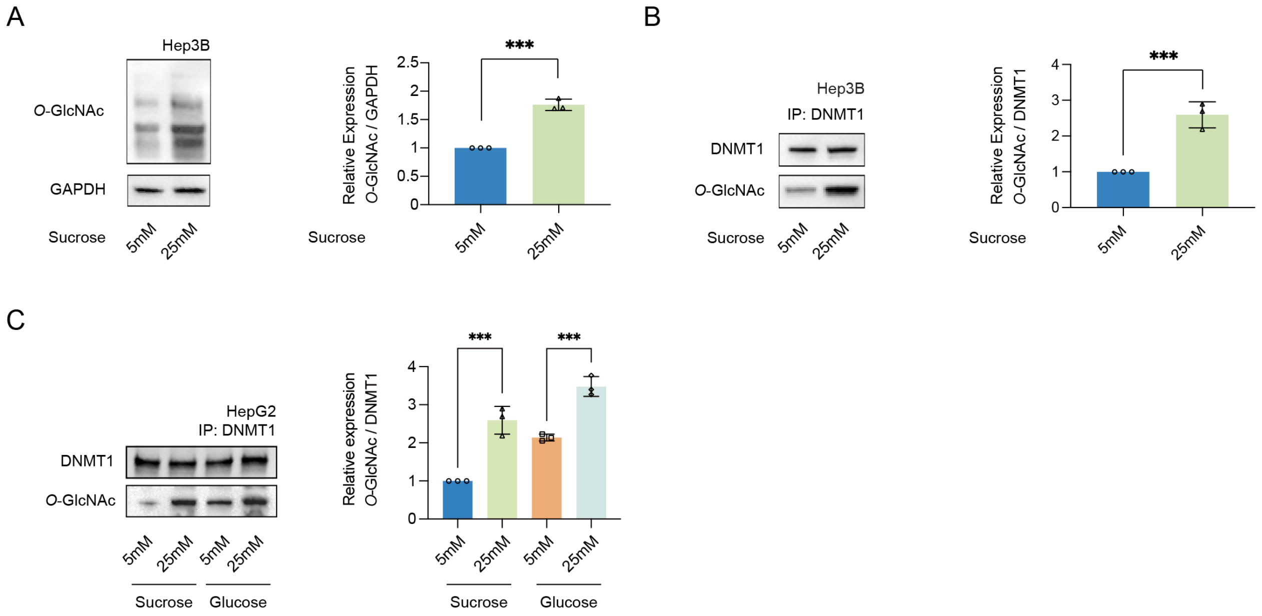

**Figure 1—figure supplement 3.** Global protein *O*-GlcNAcylation was induced with high concentrations of sucrose. (**A**) Hep3B cells were treated sucrose (5mM or 25mM). Shown are immunoblots of collected lysates using antibody targeting *O*-GlcNAc, and GAPDH (*n = 3*). (**B**) Lysates of Hep3B treated with sucrose were immunoprecipitated with DNMT1 and immunoprecipitates were immunoblotted with antibody targeting *O*-GlcNAc (n = 3). (**C**) HepG2 cells were treated 5mM glucose or sucrose, or 25mM glucose or sucrose. Lysates of HepG2 treated with glucose were immunoprecipitated with DNMT1 and immunoprecipitates were immunoblotted with antibody targeting *O*-GlcNAc (*n = 3*). ***$p < 0.0001$ by Student's *t*-test (**A-C**); Data are represented as mean ± SD from three replicates of each sample.

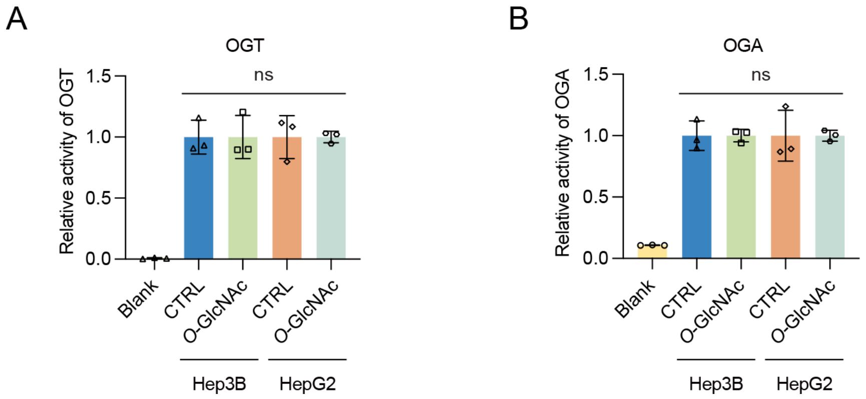

**Figure 1—figure supplement 4.** The enzymatic activity of OGT or OGA was not significantly changed by glucose treatment. (**A**) OGT activity was measured with low (5mM, CTRL) or high glucose with TMG (25mM, *O*-GlcNAc) using the UDP-Glo Glycosyltransferase activity kit (Promega) (*n* = *3*). (**B**) OGA activity was measured with low (5mM, CTRL) or high glucose with TMG (25mM, *O*-GlcNAc) using the *O*-GlcNAcase (OGA, NAG or MGEA5) assay kit (Biomedical Research Service & Clinical Application) (*n* = *3*). ns, not significant; Data are represented as mean ± SD from three replicates of each sample.

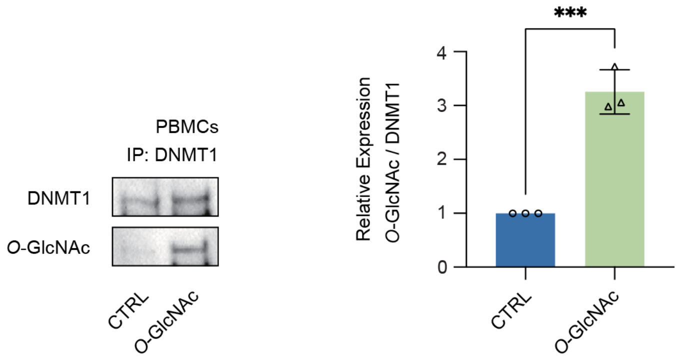

**Figure 1—figure supplement 5.** DNMT1 can be *O*-GlcNAcylated in primary cells (peripheral blood mononuclear cells, PBMCs). Pooled PBMCs treated with low (5mM, CTRL) or high glucose with TMG (25mM, *O*-GlcNAc) were immunoprecipitated with antibody targeting DNMT1 and immunoblotted for *O*-GlcNAc ($n = 3$). ***$p < 0.0001$ by Student's *t*-test; Data are represented as mean ± SD from three replicates of each sample.

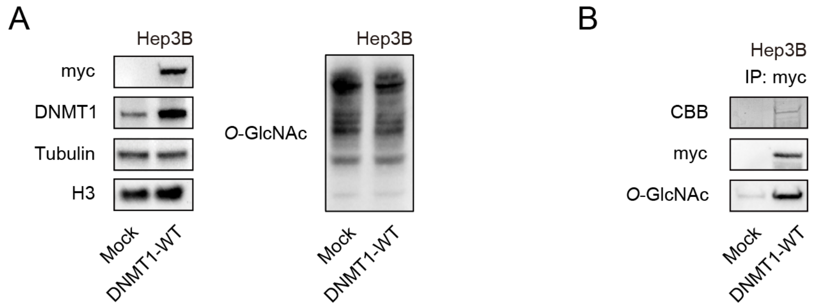

**Figure 2—figure supplement 1.** Myc-DNMT1-WT in Hep3B cells can be *O*-GlcNAcylated. (**A**) Myc-DNMT1-WT were transfected into Hep3B cells. Shown are immunoblots of treated DNMT1-WT lysates performed with antibodies targeting Myc, DNMT1, Tubulin, H3, and *O*-GlcNAc ($n$ = 3). (**B**) Lysates from treated DNMT1-WT in (**A**) were immunoprecipitated with Myc antibody. Shown are immunoblots of immunoprecipitates performed with antibodies targeting *O*-GlcNAc and CBB stained gel.

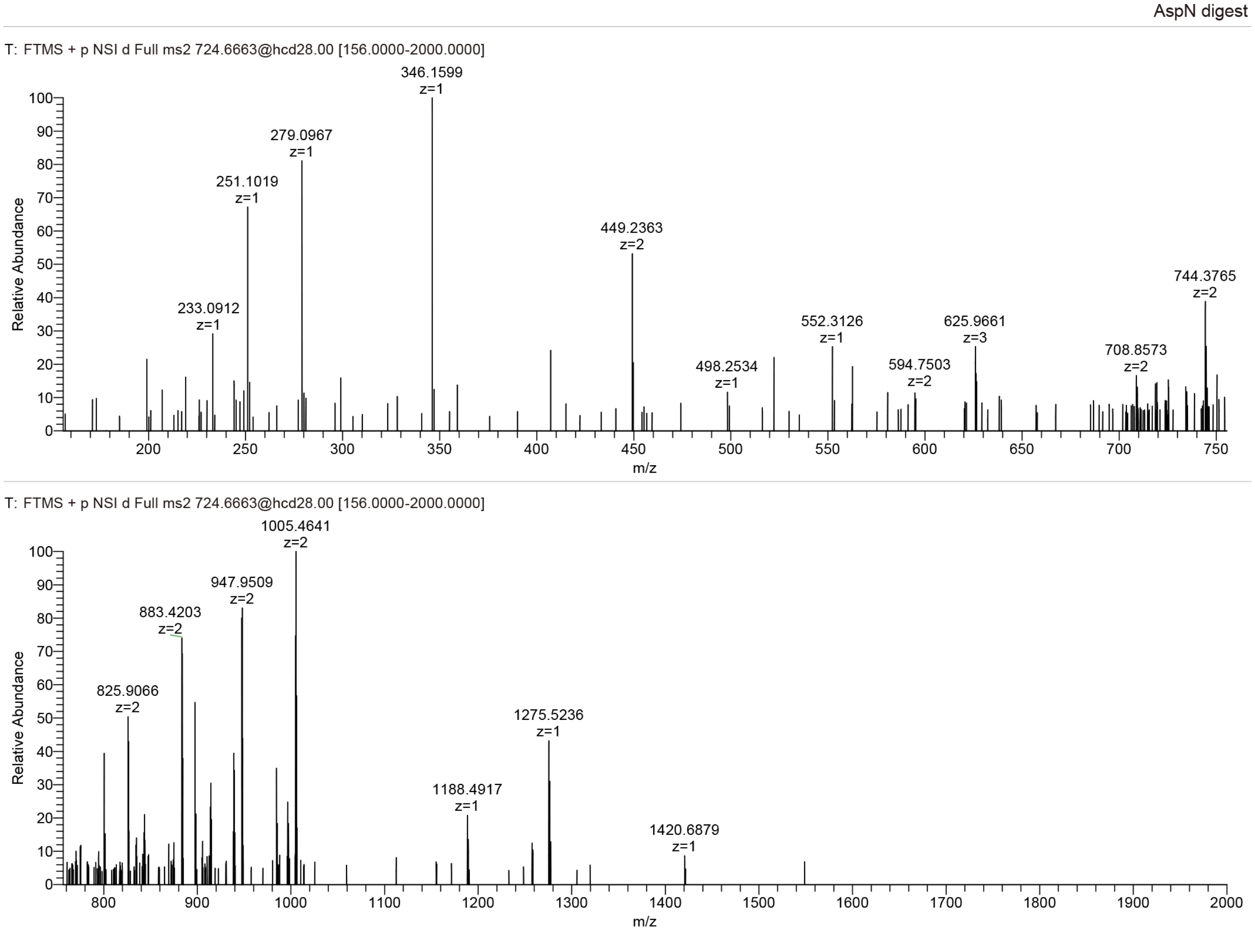

**Figure 2—figure supplement 2.** Tandem MS/MS peaks of *O*-GlcNAcylated DNMT1 peptides.

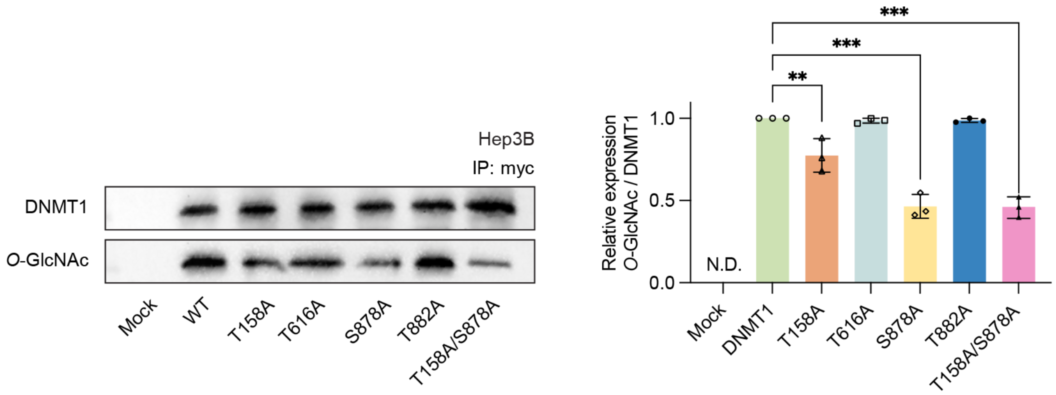

**Figure 2—figure supplement 3.** Loss of both threonine and serine at positions T158 and S878 resulted in a loss of *O*-GlcNAcylation. Each immunoprecipitated DNMT1 wild type and substituted mutants including (DNMT1-T158A/S878A double mutant) was immunoblotted with an *O*-GlcNAc antibody ($n = 3$). **$p < 0.0005$; ***$p < 0.0001$ by Student's *t*-test; N.D., not detected; Data are represented as mean ± SD from three replicates of each sample.

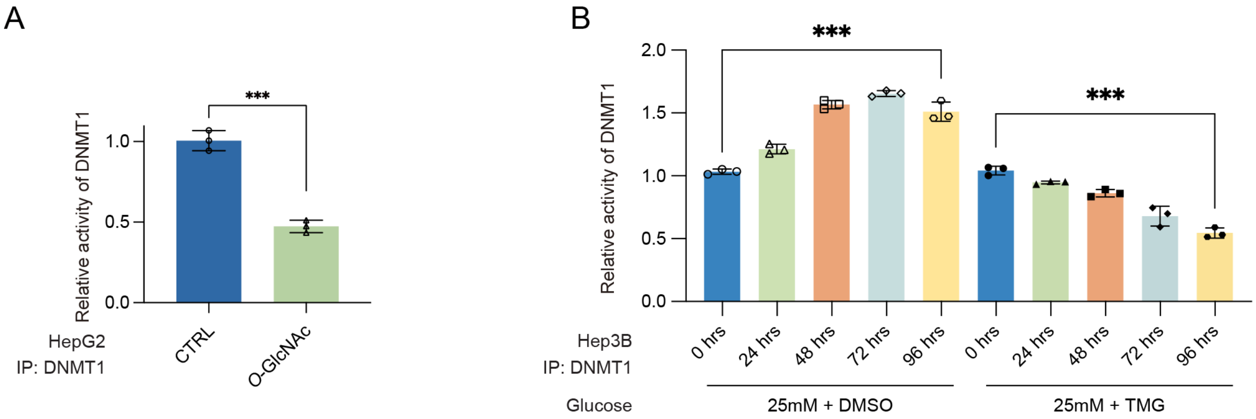

**Figure 3—figure supplement 1.** Site specific *O*-GlcNAcylation at DNMT1 sites abrogate the function of methyltransferase and DNA loss of methylation at CpG island under high glucose/TMG conditions. (**A**) HepG2 cells were treated with low (5mM, CTRL) or high glucose with TMG (25mM, *O*-GlcNAc). Shown are absorbance of DNA methyltransferase activity performed with DNA methyltransferase activity kit. (*n = 3*, technical replicates from 3 biological replicates for each strain). (**B**) Hep3B cells were treated 25mM glucose with or without Thiamet-G by time dependent. Shown are absorbance of DNA methyltransferase activity performed with DNA methyltransferase activity kit. (*n = 3*, technical replicates from 3 biological replicates for each strain). ***$p < 0.0001$ by Student's *t*-test (**A** and **B**); Data are represented as mean ± SD from three replicates of each sample.

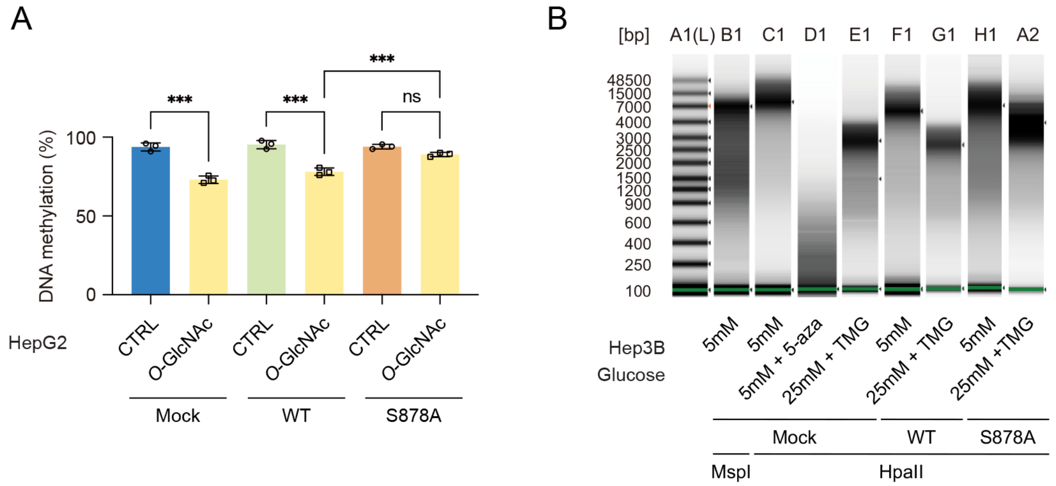

**Figure 3—figure supplement 2.** The methylation loss by high glucose/TMG conditions was not apparent in the DNMT1-S878A mutant. (**A**) Each HepG2 and Myc-DNMT1 overexpressed mutants (DNMT1-WT or DNMT1-S878A) were treated with low (5mM, CTRL) or high glucose with TMG (25mM, *O*-GlcNAc). Shown are absorbance of global DNA methylation of LINE-1 performed with global DNA methylation LINE-1 kit. (*n = 3*, technical replicates from 3 biological replicates for each strain). (**B**) DNA was extracted from Hep3B and Myc-DNMT1 overexpressed mutants (DNMT1-WT or DNMT1-S878A) were treated 5mM glucose, or 25mM glucose, or 5-aza (negative control) with MspI (negative control) or HpaII. Shown are extracted genomic DNA samples and analyze on the 4200 TapeStation System with the Genomic DNA Screen Tape assay with methylation sensitive enzyme using MspI or HpaII (*n = 3*, technical replicates from 3 biological replicates for each strain). ***$p < 0.0001$ by Student's *t*-test (**A**); ns, not significant; Data are represented as mean ± SD from three replicates of each sample.

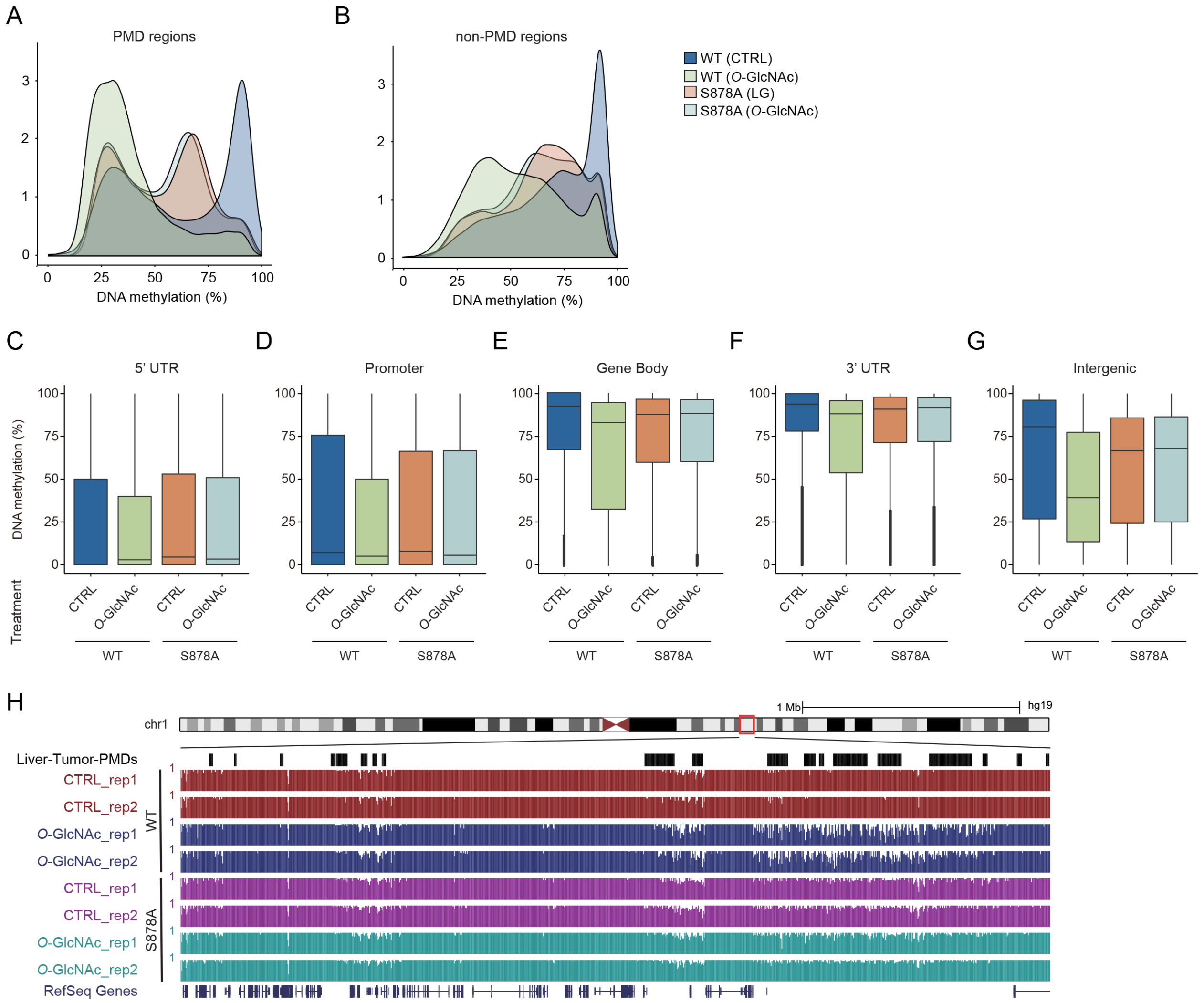

**Figure 4—figure supplement 1.** DNA loss of methylation by increased global *O*-GlcNAcylation decreases. Density plot of DNA methylation for DNMT1-WT and DNMT1-S878A cells and low or high glucose. (**A**) PMD regions, (**B**) non-PMD regions. (**C**-**G**) Bar graphs represent percentage of global DNA methylation of wild type and DNMT1 mutants (DNMT1-WT or DNMT1-S878A) which treated 5mM glucose, or 25mM glucose with Thiamet-G. (**C**) 5'UTR, (**D**) Promoter, (**E**) Gene body, (**F**) 3'UTR, (**G**) Intergenic regions. (**H**) Genome browser screenshot of DNA methylation data at a differentially methylated region by glucose concentration.

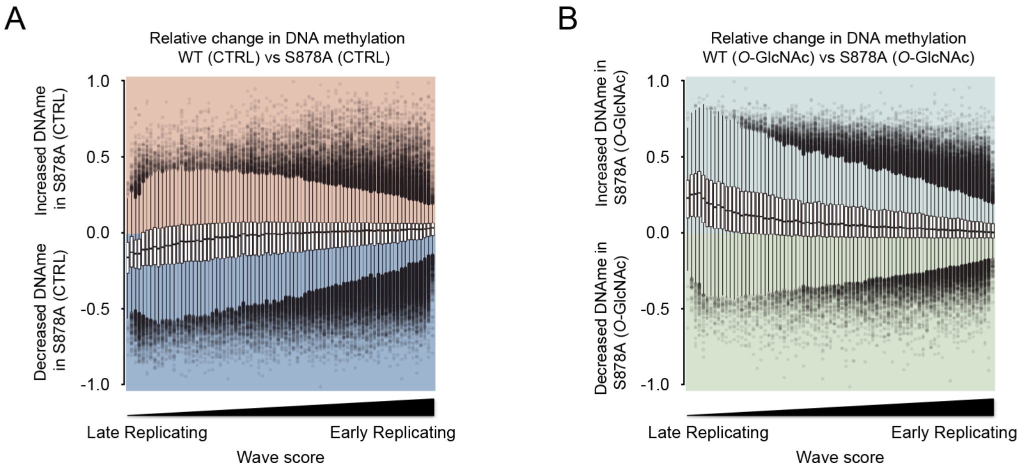

**Figure 4—figure supplement 2.** Methylation changes from *O*-GlcNAcylation of DNMT1 in DNMT1-S878A mutant. The loss of methylation was not observed in S878A mutant cells. (**A**, **B**) The distribution of each DNA methylation was divided by DNA replication timing.

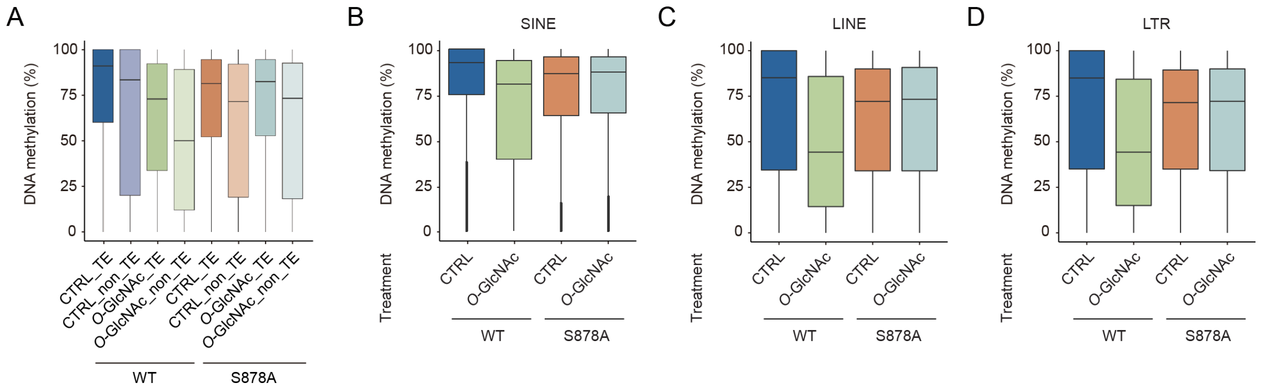

**Figure 4—figure supplement 3.** DNA loss of methylation by increased global *O*-GlcNAcylation decreases around the transposable elements (TEs) regions. (**A**) Boxplot represents the levels of DNA methylation on the TE regions or non-TE regions of each Myc-DNMT1 overexpressed mutants (DNMT1-WT or DNMT1-S878A) were treated 5mM glucose, or 25mM glucose with Thiamet-G. (**B**-**D**) bar graphs represent percentage of global DNA methylation of wild type and DNMT1 mutants (DNMT1-WT or DNMT1-S878A) which treated 5mM glucose, or 25mM glucose with Thiamet-G. (B) SINE, (C) LINE, (D) LTR regions.

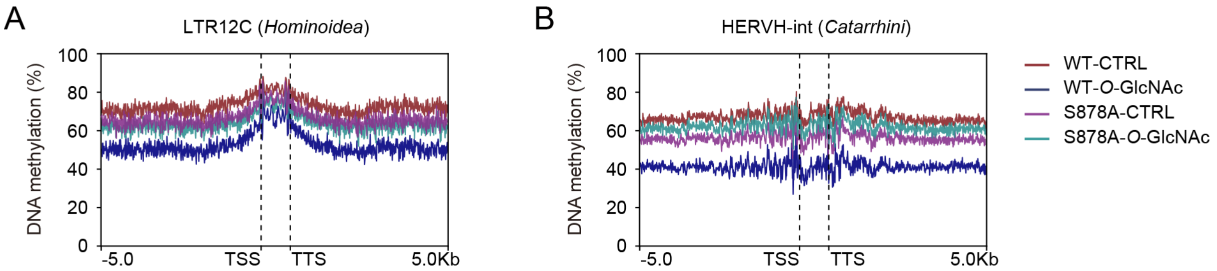

**Figure 4—figure supplement 4.** DNA loss of methylation by increased global *O*-GlcNAcylation decreases. Evolutionarily recent TEs are more likely to lose methylation than older elements in a variety of systems. Shown are methylation density around LTR12C regions of each Myc-DNMT1 overexpressed mutants (DNMT1-WT or DNMT1-S878A) were treated 5mM glucose, or 25mM glucose with Thiamet-G.

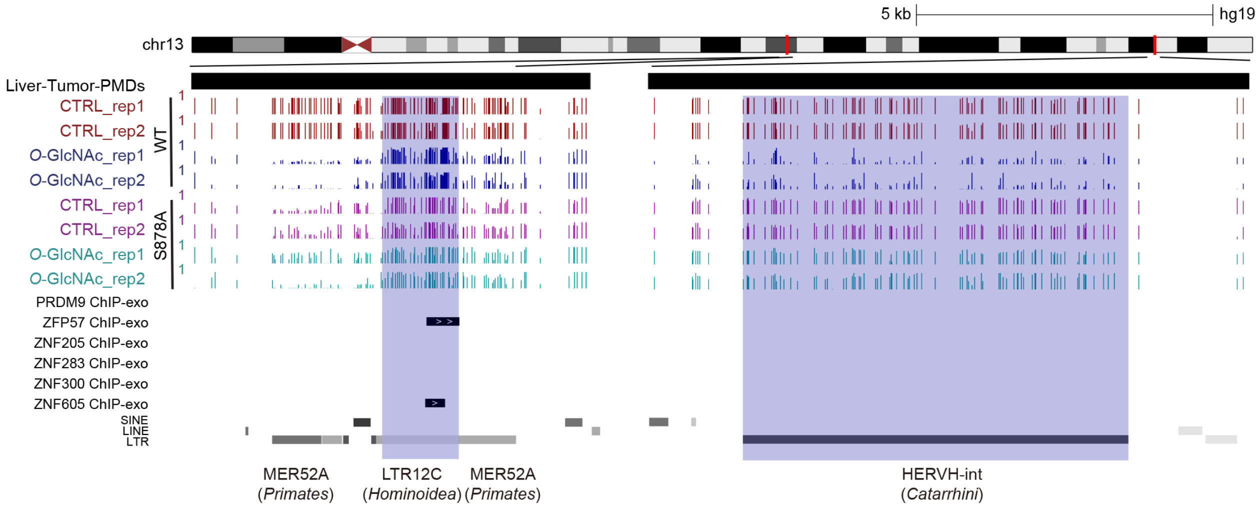

**Figure 4—figure supplement 5.** ZFP57 and ZNF605 demonstrate binding to a significant number of LTR12C elements present in liver cancer PMDs. Genome browser screenshot of DNA methylation data LTR12C elements (blue) that demonstrate binding with ZFP57 and ZNF605 by glucose concentration.

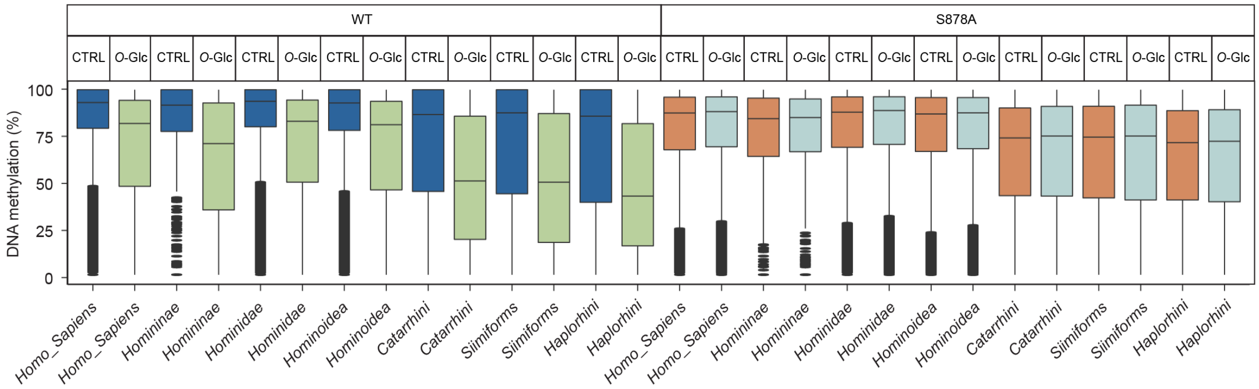

**Figure 4—figure supplement 6.** Evolutionarily recent elements are less likely to lose methylation induced by *O*-GlcNAcylation of DNMT1. Boxplot represents the DNA methylation by clades of the human genome (*Homo sapiens* to *Haplorhini*). CTRL: control; *O*-Glc: *O*-GlcNAc.

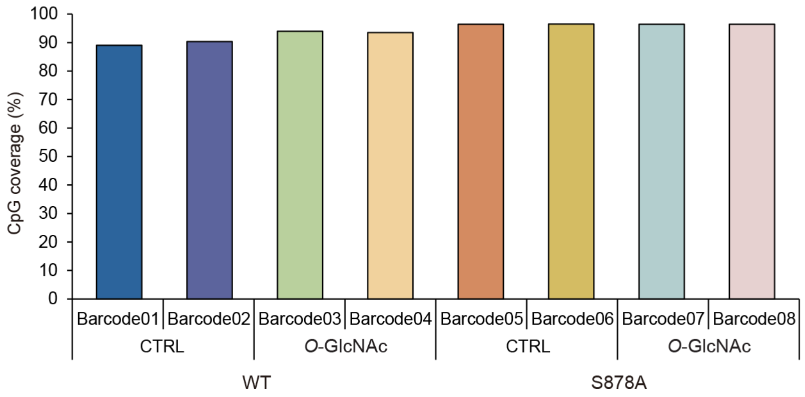

**Figure 4—figure supplement 7.** Only loci with greater than 5x coverage were retained for analysis, comprising 90% of CpGs in the genome. Shown are overall CpGs sites that detected with over 5x coverage DNA methylation analysis using Nanopore technology PromethION sequencer. Each condition is biological replicated.

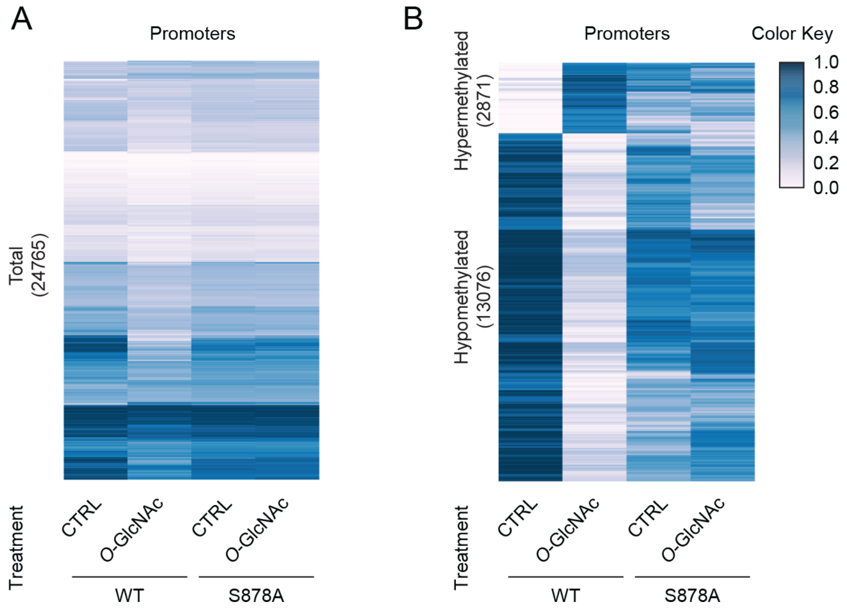

**Figure 5—figure supplement 1.** Heatmap representation of promoter DNA methylation for DNMT1-WT and DNMT1-S878A cells under with low (5mM, CTRL) or high glucose with TMG (25mM, *O*-GlcNAc) at gene poor and gene rich regions. (A) Whole promoters, (B) Differentially methylated promoters.

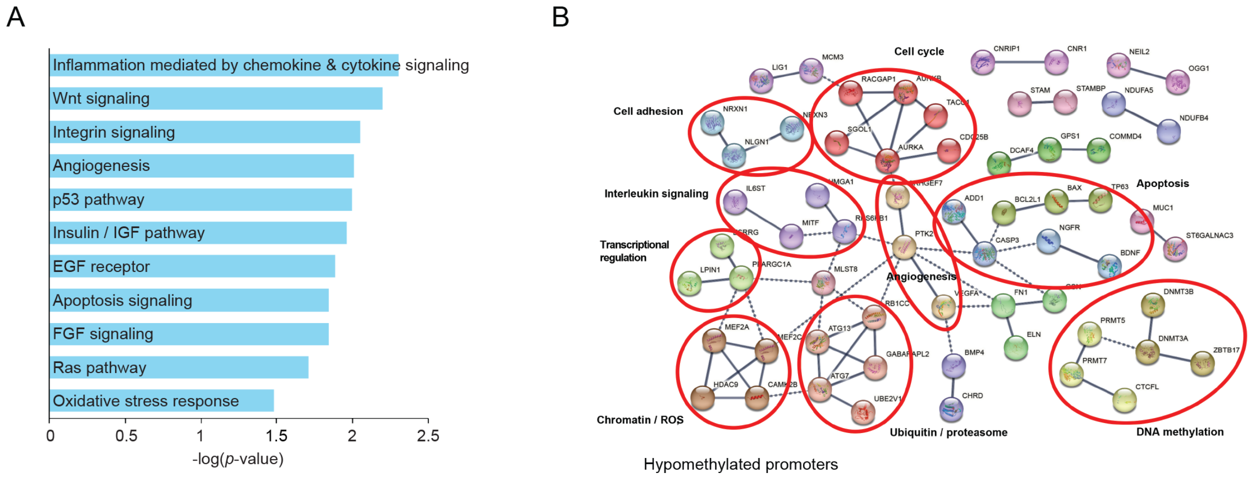

**Figure 5—figure supplement 2.** DNA loss of methylation within promoter region by increased global *O*-GlcNAcylation impact different gene pathways. (A) Gene ontology of the top 11 pathways of hypomethylated promoter DNA by high glucose treatment. (B) Gene interaction map of hypomethylated promoter DNA of DNMT1-WT by high glucose treatment.

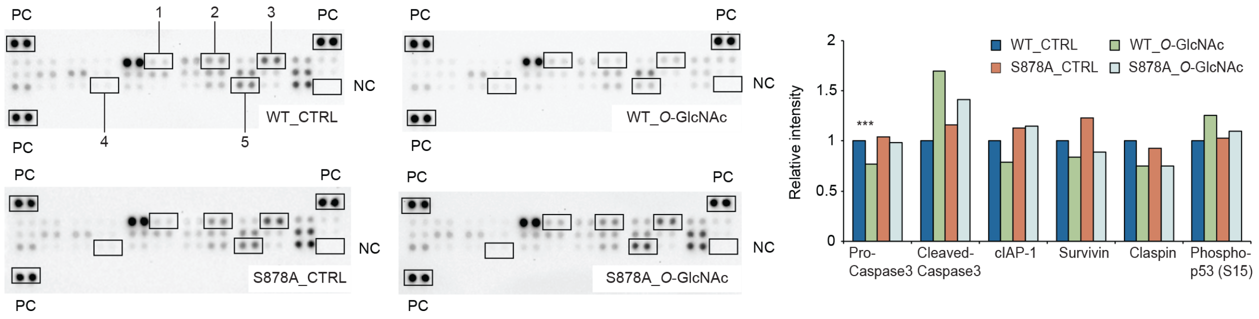

**Figure 5—figure supplement 3.** Quantitative analysis of human apoptosis related proteins in DNMT1-WT and DNMT1-S878A by high glucose treatment using Proteome profiler (*n = 3*). PC, positive control; NC, negative control; 1. cleaved-caspase3; 2, cIAP1; 3, claspin; 4, phospho-p53(S15); 5, survivin. ***$p < 0.0001$ by Student's t-test; Data are represented as mean ± SD from three replicates of each sample.

### Supplementary Tables

| Protein name | Locations | Predicted *O*-GlcNAcylated sites | Score |
|---|---|---|---|
| DNMT1 (P26358) | 5 | -MPAR T APARV | 0.8089 |
| | 158 | PSPRI T RKSTR | 0.9368 |
| | 161 | RITRK S TRQTT | 0.7442 |
| | 162 | ITRKS T RQTTI | 0.7014 |
| | 165 | KSTRQ T TITSH | 0.7822 |
| | 166 | STRQT T ITSHF | 0.7471 |
| | 168 | RQTTI T SHFAK | 0.6403 |
| | 534 | NKIET T VPPSG | 0.7764 |
| | 616 | KDRGP T KATTT | 0.8524 |
| | 801 | WFCAG T DTVLG | 0.5094 |
| | 882 | ESPPK T QPTED | 0.9432 |
| | 895 | FKFCV S CARLA | 0.6618 |
| | 977 | HYRKY S DYIKG | 0.3536 |
| | 1034 | KSTPA S YHADI | 0.7661 |
| | 1076 | CVQVY S MGGPN | 0.499 |
| | 1122 | KGKPK S QACEP | 0.7928 |

**Table S1.** Prediction of *O*-GlcNAcylated sites within DNMT1 using OGTSite.

| Gene Symbol | Accession [#] | Coverage [%] | Unique Peptides [#] | AAs [#] | MW [kDa] | calc. [pI] | Modifications |
|---|---|---|---|---|---|---|---|
| DNMT1 | P26358 | 96 | 380 | 1616 | 183.1 | 7.75 | HexNAc [S878]; Phospho [T208; S209; S394; S398; S714; S953; S954; S1105; S1122] |
| PRMT5 | O14744 | 73 | 71 | 637 | 72.6 | 6.29 | |
| PCNA | P12004 | 100 | 58 | 261 | 28.8 | 4.69 | |
| USP7 | Q93009 | 38 | 43 | 1102 | 128.2 | 5.55 | |
| HSPA1 | P0DMV9 | 60 | 42 | 641 | 70 | 5.66 | |
| HSPA8 | P11142 | 57 | 40 | 646 | 70.9 | 5.52 | |
| HSPD1 | P10809 | 57 | 34 | 573 | 61 | 5.87 | |
| RBM10 | P98175 | 36 | 33 | 930 | 103.5 | 5.97 | Phospho [S554] |
| HSPA9 | P38646 | 50 | 29 | 679 | 73.6 | 6.16 | |
| WDR77 | Q9BQA1 | 78 | 28 | 342 | 36.7 | 5.17 | Phospho [T5] |
| PPM1B | O75688 | 48 | 24 | 479 | 52.6 | 5.05 | |
| EIF4B | P23588 | 43 | 19 | 611 | 69.1 | 5.73 | Phospho [S459] |
| OTUD4 | Q01804 | 19 | 16 | 1114 | 124 | 6.71 | |
| TUBB | P07437 | 39 | 15 | 444 | 49.6 | 4.89 | |
| STK38L | Q9Y2H1 | 36 | 13 | 464 | 54 | 6.81 | Phospho [S282] |
| TRIM21 | P19474 | 33 | 13 | 475 | 54.1 | 6.38 | |
| HSPA5 | P11021 | 23 | 13 | 654 | 72.3 | 5.16 | |
| KRT10 | P13645 | 19 | 13 | 584 | 58.8 | 5.21 | |
| ERH | P84090 | 79 | 9 | 104 | 12.3 | 5.92 | |
| RPL27A | P46776 | 49 | 9 | 148 | 16.6 | 11 | |
| CLNS1A | P54105 | 44 | 9 | 237 | 26.2 | 4.11 | Phospho [S102] |
| PRPF31 | Q8WWY3 | 20 | 8 | 499 | 55.4 | 5.78 | |
| PFKFB3 | Q16875 | 16 | 8 | 520 | 59.6 | 8.21 | |
| KIF11 | P52732 | 11 | 8 | 1056 | 119.1 | 5.64 | |
| SNRPD2 | P62316 | 81 | 7 | 118 | 13.5 | 9.91 | |
| KRT1 | P04264 | 10 | 7 | 644 | 66 | 8.12 | |
| MAP1B | P46821 | 4 | 7 | 2468 | 270.5 | 4.81 | |
| MYCBP | Q99417 | 48 | 6 | 103 | 12 | 5.91 | |
| SPIN1 | Q9Y657 | 24 | 6 | 262 | 29.6 | 6.96 | |
| NPM1 | P06748 | 17 | 6 | 294 | 32.6 | 4.78 | |
| KRT2 | P35908 | 13 | 6 | 639 | 65.4 | 8 | |
| RPS11 | P62280 | 35 | 5 | 158 | 18.4 | 10.3 | |
| HNRNPK | P61978 | 17 | 5 | 463 | 50.9 | 5.54 | |
| STK38 | Q15208 | 13 | 5 | 465 | 54.2 | 7.15 | Phospho [S281] |
| KRT9 | P35527 | 12 | 5 | 623 | 62 | 5.24 | |
| RIOK1 | Q9BRS2 | 12 | 5 | 568 | 65.5 | 6.19 | |
| BCLAF1 | Q9NYF8 | 7 | 5 | 920 | 106.1 | 9.98 | Phospho [S512; S531] |
| TMPO | P42166 | 7 | 5 | 694 | 75.4 | 7.66 | |
| QPCTL | Q9NXS2 | 15 | 4 | 382 | 42.9 | 9.82 | |
| DBT | P11182 | 12 | 4 | 482 | 53.5 | 8.51 | |
| TUFM | P49411 | 11 | 4 | 452 | 49.5 | 7.61 | |
| SLC25A5 | P05141 | 11 | 4 | 298 | 32.8 | 9.69 | |
| PABPC1 | P11940 | 5 | 4 | 636 | 70.6 | 9.5 | |
| HYDIN | Q4G0P3 | 1 | 4 | 5121 | 575.5 | 6.06 | |
| RPL29 | P47914 | 25 | 3 | 159 | 17.7 | 11.66 | |
| RPS17 | P08708 | 21 | 3 | 135 | 15.5 | 9.85 | |
| EEF1B2 | P24534 | 18 | 3 | 225 | 24.7 | 4.67 | |
| C11orf84 | Q9BUA3 | 15 | 3 | 381 | 41 | 5.01 | Phospho [S308] |

| | | | | | | |
|---|---|---|---|---|---|---|
| PRDX1 | Q06830 | 15 | 3 | 199 | 22.1 | 8.13 |
| RPS2 | P15880 | 14 | 3 | 293 | 31.3 | 10.24 |
| PSMD4 | P55036 | 12 | 3 | 377 | 40.7 | 4.79 |
| PSMC2 | P35998 | 10 | 3 | 433 | 48.6 | 5.95 |
| C1QBP | Q07021 | 9 | 3 | 282 | 31.3 | 4.84 |
| RPL38 | P63173 | 36 | 2 | 70 | 8.2 | 10.1 |
| SNRPD1 | P62314 | 18 | 2 | 119 | 13.3 | 11.56 |
| RPS25 | P62851 | 17 | 2 | 125 | 13.7 | 10.11 |
| PTS | Q03393 | 16 | 2 | 145 | 16.4 | 6.68 |
| RPS12 | P25398 | 15 | 2 | 132 | 14.5 | 7.21 |
| PRDX5 | P30044 | 14 | 2 | 214 | 22.1 | 8.7 |
| RPL13 | P26373 | 14 | 2 | 211 | 24.2 | 11.65 |
| RPS3 | P23396 | 12 | 2 | 243 | 26.7 | 9.66 |
| PSMD13 | Q9UNM6 | 8 | 2 | 376 | 42.9 | 5.81 |
| CAPZA1 | P52907 | 8 | 2 | 286 | 32.9 | 5.69 |
| YWHAB | P31946 | 8 | 2 | 246 | 28.1 | 4.83 |
| RPS24 | P62847 | 8 | 2 | 133 | 15.4 | 10.78 |
| PSMC4 | P43686 | 6 | 2 | 418 | 47.3 | 5.21 |
| CMBL | Q96DG6 | 6 | 2 | 245 | 28 | 7.18 |
| PSPC1 | Q8WXF1 | 6 | 2 | 523 | 58.7 | 6.67 |
| PTBP1 | P26599 | 5 | 2 | 531 | 57.2 | 9.17 |
| RPL11 | P62913 | 5 | 2 | 178 | 20.2 | 9.6 |
| PSMC1 | P62191 | 5 | 2 | 440 | 49.2 | 6.21 |
| P4HB | P07237 | 5 | 2 | 508 | 57.1 | 4.87 |
| PSMD12 | O00232 | 5 | 2 | 456 | 52.9 | 7.65 |
| HNRNPH1 | P31943 | 4 | 2 | 449 | 49.2 | 6.3 |
| CCT5 | P48643 | 4 | 2 | 541 | 59.6 | 5.66 |
| HSPH1 | Q92598 | 3 | 2 | 858 | 96.8 | 5.39 |
| BACE2 | Q9Y5Z0 | 3 | 2 | 518 | 56.1 | 5.15 |
| ADRB2 | P07550 | 3 | 2 | 413 | 46.4 | 7.03 |
| OSBPL1A | Q9BXW6 | 3 | 2 | 950 | 108.4 | 6.38 |
| FAM160A1 | Q05DH4 | 3 | 2 | 1040 | 116.5 | 4.86 |
| HSPA4L | O95757 | 3 | 2 | 839 | 94.5 | 5.88 |
| RNF219 | Q5W0B1 | 3 | 2 | 726 | 81.1 | 5.72 |
| THRAP3 | Q9Y2W1 | 2 | 2 | 955 | 108.6 | 10.15 |
| ITPRIP | Q8IWB1 | 2 | 2 | 547 | 62 | 5.88 |
| SPEF2 | Q9C093 | 1 | 2 | 1822 | 209.7 | 5.54 |
| ERCC6L | Q2NKX8 | 1 | 2 | 1250 | 141 | 5.31 |
| SRCAP | Q6ZRS2 | 1 | 2 | 3230 | 343.3 | 5.96 |

**Table S2.** List of total identified proteins.

| | | |
|---|---|---|
| **Antibodies** | | |
| Alpha-tubulin (11H10) | Cell Signaling Technology | 2125S |
| DNMT1 (60B1220.1) | Novus Biologicals | NB100-56519 |
| DNMT1 (H-12) | Santa Cruz Biotechnology | sc-271729 |
| GAPDH | Abcam | ab181602 |
| H3 | | ab1791 |
| Myc tag [Myc.A7] | | ab18185 |
| *O*-GlcNAc (RL2) | | ab2739 |
| OGT | | ab177941 |
| **Chemicals** | | |
| OSMI-4 | Selleck Chem | S8910 |
| Thiamet-G | Cayman Chemical | 13237 |
| **Deposited data** | | |
| DNMT1-WT-CTRL | This paper | GEO: GSE201470 |
| DNMT1-WT-*O*-GlcNAc | This paper | GEO: GSE201470 |
| DNMT1-S878A-CTRL | This paper | GEO: GSE201470 |
| DNMT1-S878A-*O*-GlcNAc | This paper | GEO: GSE201470 |
| Partially methylated domains of liver cancer | Li et al., 2016 | GEO: GSE70091 |
| RNA sequencing | Chang et al., 2014 | GEO: GSE49994 |
| **Oligonucleotides** | | |
| DNMT1-T158A | (Forward): | 5'-agccccaggatt CGA aggaaaagcacc-3' |
| | (Reverse): | 5'-ggtgcttttcct TCG aatcctggggct-3' |
| DNMT1-T616A | (Forward): | 5'-gacaggggaccc GCG aaagccaccacc-3' |
| | (Reverse): | 5'-ggtggtggcttt CGC gggtcccctgtc-3' |
| DNMT1-S878A | (Forward): | 5'-gcgagattcgag GAG cctccaaaaacc-3' |
| | (Reverse): | 5'-ggtttttggagg CTC ctcgaatctcgc-3' |
| DNMT1-S878D | (Forward): | 5'-gcgagattcgag GAC cctccaaaaacc-3' |
| | (Reverse): | 5'-ggtttttggagg GTC ctcgaatctcgc-3' |
| DNMT1-T882A | (Forward): | 5'-tcccctccaaaa GCC cagccaacagag-3' |
| | (Reverse): | 5'-ctctgttggctg GGC ttttggaggga-3' |
| **Software and algorithms** | | |
| bedGraphToBigWig | Kent et al., 2010 | https://www.encodeproject.org/software/bedgraphtobigwig/ |
| Graphpad Prism 9 (v9.3.1) | Graphpad | https://www.graphpad.com/scientific-software/prism/ |
| Minimap2 (v2.17) (RRID:SCR_018550) | Li, 2018 | https://github.com/lh3/minimap2 |
| Nanopolish (v0.11.1) (RRID:SCR_016157) | Loman et al., 2015 | https://github.com/jts/nanopolish |
| Python (v3.8.2) | Python Core Team | https://www.python.org/ |
| R (v3.4.3) | R Core Team | https://www.r-project.org/ |
| Samtools (v1.10) (RRID:SCR_002105) | Li et al., 2009 | |

**Table S3.** List of antibodies and reagent used in this study.

**Figure 1—source data 1.** Uncropped blot files of Figure 1A-E.

**Figure 1—figure supplement 1-source data 1.** Uncropped blot files of Figure 1—figure supplement 1A and B.

**Figure 1—figure supplement 2-source data 1.** Uncropped blot files of Figure 1—figure supplement 2A and B.

**Figure 1—figure supplement 3-source data 1.** Uncropped blot files of Figure 1—figure supplement 3A-C.

**Figure 1—figure supplement 5-source data 1.** Uncropped blot files of Figure 1—figure supplement 5.

**Figure 2—source data 1.** Uncropped blot files of Figure 2C.

**Figure 2—figure supplement 1-source data 1.** Figure 2—figure supplement 1A and B.

**Figure 2—figure supplement 3-source data 1.** Figure 2—figure supplement 3.

**Figure 5—source data 1.** Raw fluorescence image files of Figure 5A and C.

**Figure 5—figure supplement 3-source data 1.** Uncropped blot files of Figure 5—figure supplement 3.
